# Comparative anatomy and phylogenetic contribution of intracranial osseous canals and cavities in armadillos and glyptodonts (Xenarthra, Cingulata)

**DOI:** 10.1101/2021.04.21.440734

**Authors:** Kévin Le Verger, Laureano R. González Ruiz, Guillaume Billet

## Abstract

The evolutionary history of the Cingulata, as for many groups, remains a highly debated topic to this day, particularly for one of their most emblematic representatives: the glyptodonts. There is no consensus among morphological and molecular phylogenies relative to their position within Cingulata. As demonstrated by recent works, the study of the internal anatomy constitutes a promising path for enriching morphological matrices for the phylogenetic study of armadillos. However, internal cranial anatomy remains under-studied in the Cingulata. Here we explored and compared the anatomy of intracranial osseous canals and cavities in a diverse sample of extant and extinct cingulates, including the earliest well-preserved glyptodont crania. The virtual 3D reconstruction (using X-ray microtomography) of selected canals, i.e., the nasolacrimal canal, the palatine canal, the sphenopalatine canal, the canal for the frontal diploic vein, the transverse canal, the orbitotemporal canal, the canal for the capsuloparietal emissary vein and the posttemporal canal, and alveolar cavities related to cranial vascularization, innervation or tooth insertion allowed us to compare the locations, trajectories and shape of these structures and to discuss their potential interest for cingulate systematics. We tentatively reconstructed evolutionary scenarios for eight selected traits on these structures, in which glyptodonts often showed a greater resemblance to pampatheres, to the genus *Proeutatus* and/or to chlamyphorines. This latter pattern was partly congruent with recent molecular hypotheses, but more research is needed on these resemblances and on the potential effects of development and allometry on the observed variations. Overall, these comparisons have enabled us to highlight new anatomical variation that may be of great interest to further explore the evolutionary history of cingulates and the origins of glyptodonts on a morphological basis.

## 1. INTRODUCTION

The peculiar morphology of xenarthrans, which include armadillos (Cingulata), sloths (Folivora) and anteaters (Vermilingua), has aroused the curiosity of anatomists since the end of the 18th century (Cuvier, 1798). Compared to other mammals, the sum of their unique characteristics has led them to be considered by some earlier workers the most unusual mammals (Vizcaíno & Loughbry, 2008; for a summary see Superina & Loughry, 2015). They represent the only known mammals with a carapace formed by a mosaic of osteoderms (fixed and mobile) covering the head, body and tail (Vizcaíno & Loughry, 2008). Cingulates include extant and extinct armadillos, among them the four extant subfamilies Dasypodinae, Euphractinae, Tolypeutinae, and Chlamyphorinae, as well as the extinct glyptodonts, a group of herbivorous and large bodied mammals appearing in the late Eocene and disappearing at the beginning of the Holocene (Delsuc et al., 2016; see also Gaudin & Lyon, 2017 for an alternative classification). The phylogenetic position of glyptodonts within Cingulata is particularly difficult to resolve based on morphological data because of their highly specialized anatomy and the large morphological gap separating them from extant armadillos (Burmeister, 1874; Carlini & Zurita, 2010; Fariña et al., 2013). This difficulty was recently emphasized by ancient DNA studies showing that phylogenetic analyses of their mitogenomes support a position for glyptodonts at odds with that proposed in prior morphological studies (Engelmann, 1985; Gaudin & Wible, 2006; Billet et al., 2011; Delsuc et al., 2016; Mitchell et al., 2016; Herrera et al., 2017).

While it has long been accepted that glyptodonts were closely related to extant armadillos (Huxley, 1864), their exact relationships with the latter were never clearly resolved on the basis morphological data. Flower (1882) initially proposed that glyptodonts were the ancestors of extant armadillos but Ameghino (1884, 1889) disputed this hypothesis, arguing that their great dental and postcranial complexity, as described by Owen (1839) was evidence of their derived nature. The nested position of glyptodonts among extant armadillos has been supported by many authors since then, including by recent phylogenetic analyses using morphological data (Castellanos, 1932, 1959; Hoffstetter, 1958; Patterson & Pascual, 1972; Fernicola, 2008; Gaudin & Wible, 2006; Billet *et al*., 2011; Herrera et al., 2017). However, the consensus stops here. Engelmann (1985) proposed a close relationship of glyptodonts with the extinct Eutatini (*Proeutatus* and *Eutatus*) and pampatheres, whereas the analysis of Gaudin & Wible (2006) retrieved glyptodonts in an apical position nested within a large clade gathering euphractines, chlamyphorines, the extinct pampatheres and some extinct eutatines. A similar hypothesis was also supported by other studies, but with much variation in the composition of this large apical clade (Fernicola, 2008; Porpino et al., 2010; Billet et al., 2011; Fernicola et al., 2017; Herrera et al., 2017). Most of these studies, however, unvaryingly proposed a close relationship between the extinct giant pampatheres and glyptodonts, in agreement with Patterson & Pascual (1972). Thanks to the progress made in ancient DNA studies and the recent extinction of glyptodonts – at the beginning of the Holocene (Messineo & Politis, 2009) – the complete mitochondrial genome of the Pleistocene glyptodont *Doedicurus* Burmeister, 1874 could be successfully assembled, which gave rise to a completely new phylogenetic hypothesis (Delsuc et al., 2016; Mitchell et al., 2016). Based on analyses of their mitogenomes, glyptodonts are nested within extant armadillos and represent the sister group of a clade formed by chlamyphorines and tolypeutines (Delsuc et al., 2016; Mitchell et al., 2016). Although analyses of morphological matrices using a molecular backbone constraint can detect morphological characters congruent with the molecular pattern (Mitchell et al., 2016), the high level of disagreement between the two data partitions may call into question the phylogenetic signal provided by morphological matrices on cingulates.

External cranial and postcranial skeletal anatomy are already well known for many groups within Cingulata, but the internal cranial anatomy remains understudied. However, its study has already provided data of systematic interest for both the extant and extinct xenarthran diversity (Zurita et al., 2011; Fernicola et al., 2012; Billet et al., 2015; Tambusso & Fariña, 2015a; Tambusso & Fariña, 2015b; Billet et al., 2017; Boscaini et al., 2018; Boscaini et al., 2020; Tambusso et al., 2021). Intracranial osseous canals provide important pathways for innervation and vascularization of the head (Evans & de Lahunta, 2012). The diversity and phylogenetic signal of their intracranial trajectories are poorly known, as these hidden structures are rarely described in mammals in general, including xenarthrans (e.g., Wible & Gaudin, 2004). In contrast, their external openings on the cranium are often described and scored in phylogenetic matrices (*e.g.*, 37/131 of cranial characters – Gaudin & Wible, 2006).

Here we present a comparative investigation of intracranial osseous canals and cavities in a diverse sample of extant and extinct cingulates, including the earliest well-preserved glyptodont crania, using X-ray microtomography. The 3D virtual reconstruction of selected canals and cavities related to cranial vascularization, innervation or upper tooth insertion enabled us to compare the locations, trajectories and shape of each homologous structure and discuss their potential interest for cingulate systematics. We tentatively reconstructed the evolutionary history of these traits using a molecularly constrained phylogeny, and we discuss the potential effects of development and allometry on their variation. These comparisons allowed us to propose new potential characters that may be used to further explore the origins of glyptodonts within cingulates on a morphological basis.

## 2. MATERIALS AND METHODS

### 2.1 Sampling

We examined the crania of 33 extant and extinct xenarthran specimens. Two specimens belonging to the Pilosa (sloths and anteaters) were chosen as outgroups. The remaining 31 crania belonged to Cingulata (Table 1). This sample included specimens representing the 9 extant genera, as well as three small developmental series in phylogenetically distant species (*Dasypus novemcinctus*, *Zaedyus pichiy*, *Cabassous unicinctus*), with the aim of better understanding the ontogenetic variation of the selected anatomical structures. The cingulate specimens also included 14 specimens belonging to fossil species, among them six glyptodonts (Table 1). This sample covered more than 40 million years of cingulate evolutionary history and includes all the major cingulate subfamilies. Our sample does not allow us to evaluate intraspecific variation in all taxa. However, we were able to estimate this variation in three species for which we dispose developmental series. For each of them, we analyzed scans of 4 additional adult specimens. The adult intraspecific sampling is available in Table S1. Species identification was based on collection data, geographical origin, cranial anatomy, and the literature on cingulate taxonomy (e.g., Scott, 1903; McBee & Baker, 1982; Wetzel, 1985; Gaudin & Wible, 2006; Wetzel et al., 2007; Hayssen, 2014; Abba et al., 2015; Carlini et al., 2016; Gaudin & Lyon, 2017; Smith & Owen, 2017). Digital data of all specimens were acquired using X-ray micro-computed tomography (μCT). Specimens were scanned on X-ray tomography imagery platforms at the American Museum of Natural History (New York, USA); the Museum national d’Histoire naturelle (France) in Paris (AST-RX platform), the University of Montpellier (France – MRI platform) and the Museum für Naturkunde (ZMB) in Berlin (Phoenix nanotom (General Electric GmbH Wunstorf, Germany)). Three-dimensional reconstructions of the selected structures were performed using stacks of digital μCT images with MIMICS v. 21.0 software (3D Medical Image Processing Software, Materialize, Leuven, Belgium). The visualization of 3D models was also conducted with AVIZO v. 9.7.0 software (Visualization Sciences Group, Burlington, MA, USA). A large part of specimens scanned will be available on MorphoMuseum for 3D models and on MorphoSource for scans. Specimens scanned by LRG will be available upon request to LRG. The list of specimens is given in Table S2.

**TABLE 1.** List of specimens. Symbol: †, extinct species; *, perinatal stage; **, juvenile; ***, subadult; ****, adult; (i), author of the genus.

| Family/Subfamily | Species | Institutional number | Locality | Period | References |
| --- | --- | --- | --- | --- | --- |
| Bradypodidae | *Bradypus tridactylus* | MNHN ZM-MO-1999-1065 | French Guiana; Petit saut rive droite | Extant | Linnaeus, 1758 |
| Myrmecophagidae | *Tamandua tetradactyla* | NHM 3-7-7-135 | Brazil; Mato Grosso; Chapada | Extant | Linnaeus, 1758 |
| Peltephilidae | *Peltephilus pumilus*† | YPM-VPPU 15391 | Argentina; Santa Cruz; Santa Cruz Formation; Coy Inlet | Early Miocene | Ameghino, 1887 |
| Dasypodinae | *Dasypus novemcinctus* * | AMNH 33150 | Colombia; Quindui; Salento | Extant | Linnaeus, 1758 |
| Dasypodinae | *Dasypus novemcinctus* ** | AMNH 133261 | Brazil; Goias; Anapolis | Extant | Linnaeus, 1758 |
| Dasypodinae | *Dasypus novemcinctus* *** | AMNH 133328 | Brazil; Mato Grosso do Sul; Maracaju | Extant | Linnaeus, 1758 |
| Dasypodinae | *Dasypus novemcinctus* **** | AMNH 255866 | Bolivia; Beni; Cercado | Extant | Linnaeus, 1758 |
| Dasypodidae | *Stegotherium tauberi*† | YPM-VPPU 15565 (type) | Argentina; Santa Cruz; Santa Cruz Formation; Killik Aike Norte | Early Miocene | González y Scillato-Yané, 2008 |
| Euphractinae | *Prozaedyus exilis*† | YPM-VPPU 15579 | Argentina; Santa Cruz; Santa Cruz Formation; Killik Aike Norte | Early Miocene | Ameghino, 1887 |
| Euphractinae | *Prozaedyus proximus*† | YPM-VPPU 15567 | Argentina; Santa Cruz; Santa Cruz Formation; Coy Inlet | Early Miocene | Ameghino, 1887 |
| Euphractinae | *Euphractus sexcinctus* | AMNH 133304 | Brazil; Goias; Anapolis | Extant | Linnaeus, 1758 |
| Euphractinae | *Chaetophractus villosus* | AMNH 173546 | South America; No more precision | Extant | Desmarest, 1804 |
| Euphractinae | *Zaedyus pichiy* * | ZMB 49039 | Argentina; Oso Marino | Extant | Desmarest, 1804 |
| Euphractinae | *Zaedyus pichiy* ** | MHNG 1627.053 | Argentina; Chubut; Punta Ninfas | Extant | Desmarest, 1804 |
| Euphractinae | *Zaedyus pichiy* *** | MHNG 1276.076 | Argentina; Chubut; Fofó Cahuel | Extant | Desmarest, 1804 |
| Euphractinae | *Zaedyus pichiy* **** | FMNH 23810 | Argentina; Chubut; No more precision | Extant | Desmarest, 1804 |
| Euphractinae | *Doellotatus* sp.† | UF 260533 | Bolivia; unnamed formation; Potosí; Quebrada Honda | Middle Miocene | Bordas, 1932 (i) |
| Tolypeutinae | *Cabassous unicinctus* ** | NBC 26326.B | Suriname; No more precision | Extant | Linnaeus, 1758 |
| Tolypeutinae | *Cabassous unicinctus* *** | AMNH 133386 | Brazil; Mato Grosso do Sul; Maracaju | Extant | Linnaeus, 1758 |
| Tolypeutinae | *Cabassous unicinctus* **** | MNHN 1999-1044 | French Guiana; No more precision | Extant | Linnaeus, 1758 |
| Tolypeutinae | *Tolypeutes matacus* | FMNH 28345 | Brazil; Mato Grosso; Descalvado | Extant | Desmarest, 1804 |
| Tolypeutinae | *Priodontes maximus* | AMNH 208104 | Zoo; No more precision | Extant | Kerr, 1792 |
| Chlamyphorinae | *Chlamyphorus truncatus* | AMNH 5487 | Argentina; Mendoza; No more precision | Extant | Harlan, 1825 |
| Chlamyphorinae | *Calyptophractus retusus* | NMNH 283134 | Bolivia; Santa Cruz; No more precision | Extant | Burmeister, 1863 |
| Eutatini | *Proeutatus lagena*† | YPM-VPPU 15613 | Argentina; Santa Cruz; Santa Cruz Formation; Coy Inlet | Early Miocene | Ameghino, 1887 |
| Pampatheriinae | *Vassallia maxima*† | FMNH P14424 | Argentina; Catamarca; Andalhuala Formation; Puerta del Corral Quemado | Late Miocene | Castellanos, 1946 |
| Glyptodontinae | *Propalaeohoplophorus minus*† | AMNH-FM 9197 | Argentina; Santa Cruz; Santa Cruz Formation; Cañadón de las Vacas | Early Miocene | Ameghino, 1891 |
| Glyptodontinae | *"Cochlops" debilis*† | YPM-VPPU 15592 | Argentina; Santa Cruz; Santa Cruz Formation; Cañadón Silva | Early Miocene | Ameghino, 1891 |
| Glyptodontinae | *Eucinepeltus complicatus*† | AMNH-FM 9248 (type) | Argentina; Santa Cruz; Santa Cruz Formation; Killik Aike Norte | Early Miocene | Brown, 1903 |
| Glyptodontinae | *"Metopotoxus" anceps*† | YPM-VPPU 15612 | Argentina; Santa Cruz; Santa Cruz Formation; Lago Pueyrredón | Early Miocene | Scott, 1903 |
| Glyptodontinae | *Glyptodon* sp.† | MNHN PAM-759 | Argentina; Buenos Aires; No more precision | Pleistocene | Owen, 1839 (i) |
| Glyptodontinae | *Glyptodon* sp.† | MNHN PAM-760 | Argentina; Buenos Aires; No more precision | Pleistocene | Owen, 1839 (i) |

**TABLE 2.**
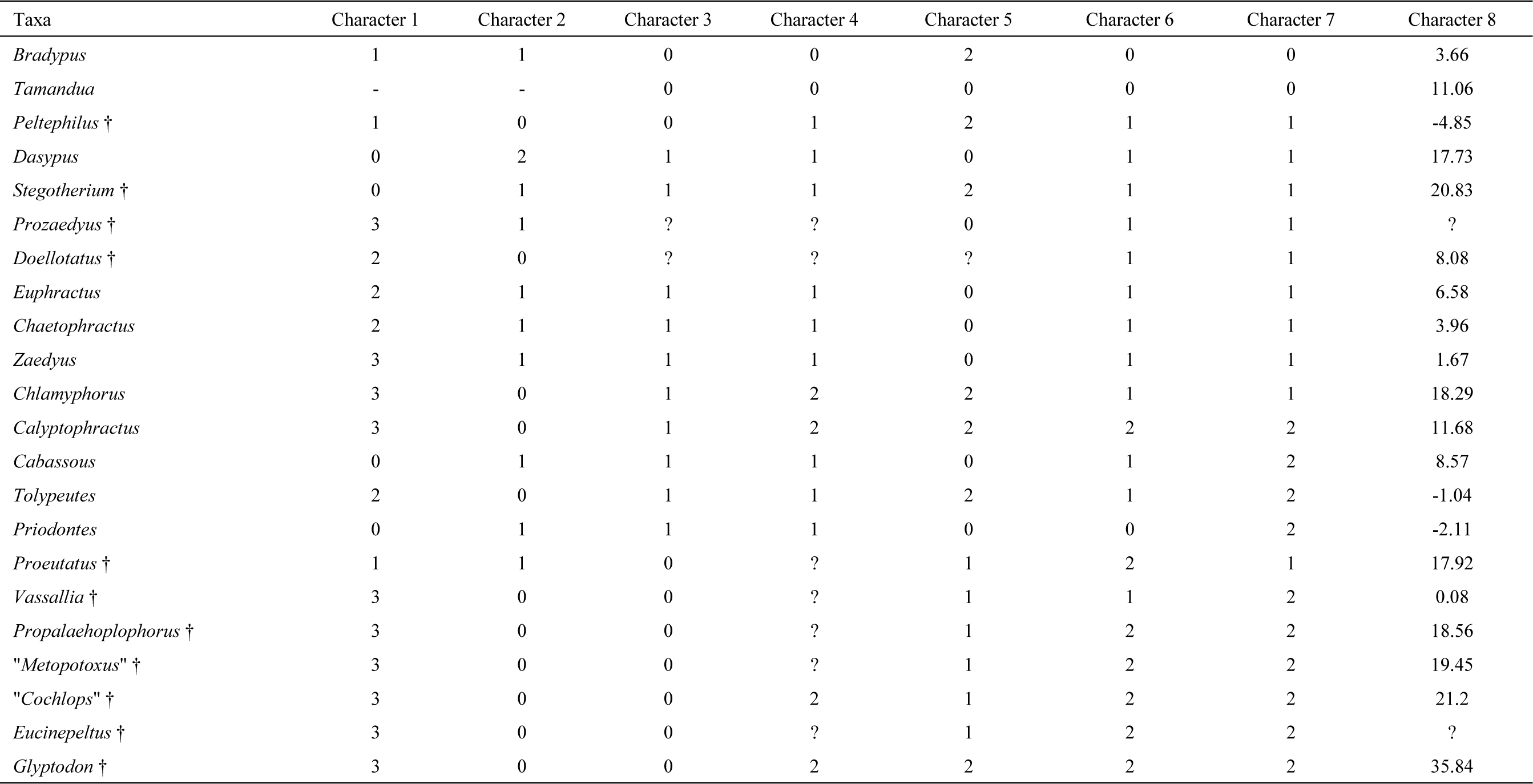
Data matrix with character scores for each genus. Character 1 and 7 are discrete and character 8 is continuous (see text). Symbols: -, not applicable; ?, missing data; †, extinct species.

### 2.2 Selected regions of interest and anatomical nomenclature

We selected several osseous anatomical complexes in the internal cranial anatomy of cingulates that are poorly known and of interest for this study such as dental alveoli and several specific intraosseous canals. The latter mostly correspond to vascular pathways and/or the courses of cranial nerves involved in the innervation of various cranial areas: the nasolacrimal canal, the palatine canal, the sphenopalatine canal, the canal for the frontal diploic vein, the transverse canal, the orbitotemporal canal, the canal for the capsuloparietal emissary vein and the posttemporal canal.

Our study is focused only on the cranium and does not include the mandible. Therefore, we are only interested in the upper dentition. Except for *Dasypus*, which represents a special case (see Ciancio et al., 2012), the homologies in the dental row between the different species of Cingulata are not known mainly because of the drastic reduction in dental complexity (Vizcaíno, 2009). For glyptodonts, previous studies conventionally designated the whole set of teeth as molariform but without further precision (e.g., González Ruiz et al., 2015). We therefore chose to treat all teeth as molariform (= Mf; see Herrera et al., 2017 for a different nomenclature and Table S3 for the dental formula of each specimen of our study with greater precision).

Only those intracranial canals whose sampled variation appeared to bear clear systematic information were selected in this work (Figure 1). Non-selected canals generally showed asymmetric and intraspecific variation according to our intraspecific sample (Table S1 – e.g., hypoglossal canal; trajectory of the internal carotid artery – see also Patterson et al., 1989; Gaudin, 1995) or provided no new data (e.g., infraorbital canal) with respect to what was already described or scored for the systematics of the group (Wible & Gaudin, 2004; Gaudin & Wible, 2006; Gaudin & Lyon, 2017). Other non-selected canals were rarely visible in all specimens (often due to taphonomy – not in place and/or obscured by a hard and dense matrix) and were therefore difficult to compare. Specimens that presented a canal that was too incomplete are not described in the relevant section. For the identification and nomenclature of the selected intracranial canals, our study used previous work describing intracranial anatomy in cingulates and eutherians in general (Thewissen 1989; Wible, 1993; Gaudin, 2004; Wible & Gaudin 2004; Evans & de Lahunta, 2012; Muizon et al., 2015; Gaudin & Lyon, 2017). We have indicated for each selected region of interest: the variation of these regions during ontogeny, a synthetic comparison among specimens, and the formalization of potential discrete or continuous characters to highlight potential evolutionary scenarios to be mapped onto the tree of cingulates. The formalization of new characters was performed based on observations that were stable among glyptodonts and were shared with some non-glyptodonts cingulates, so that they could provide pertinent information when investigating glyptodont origins.

**FIGURE 1.**
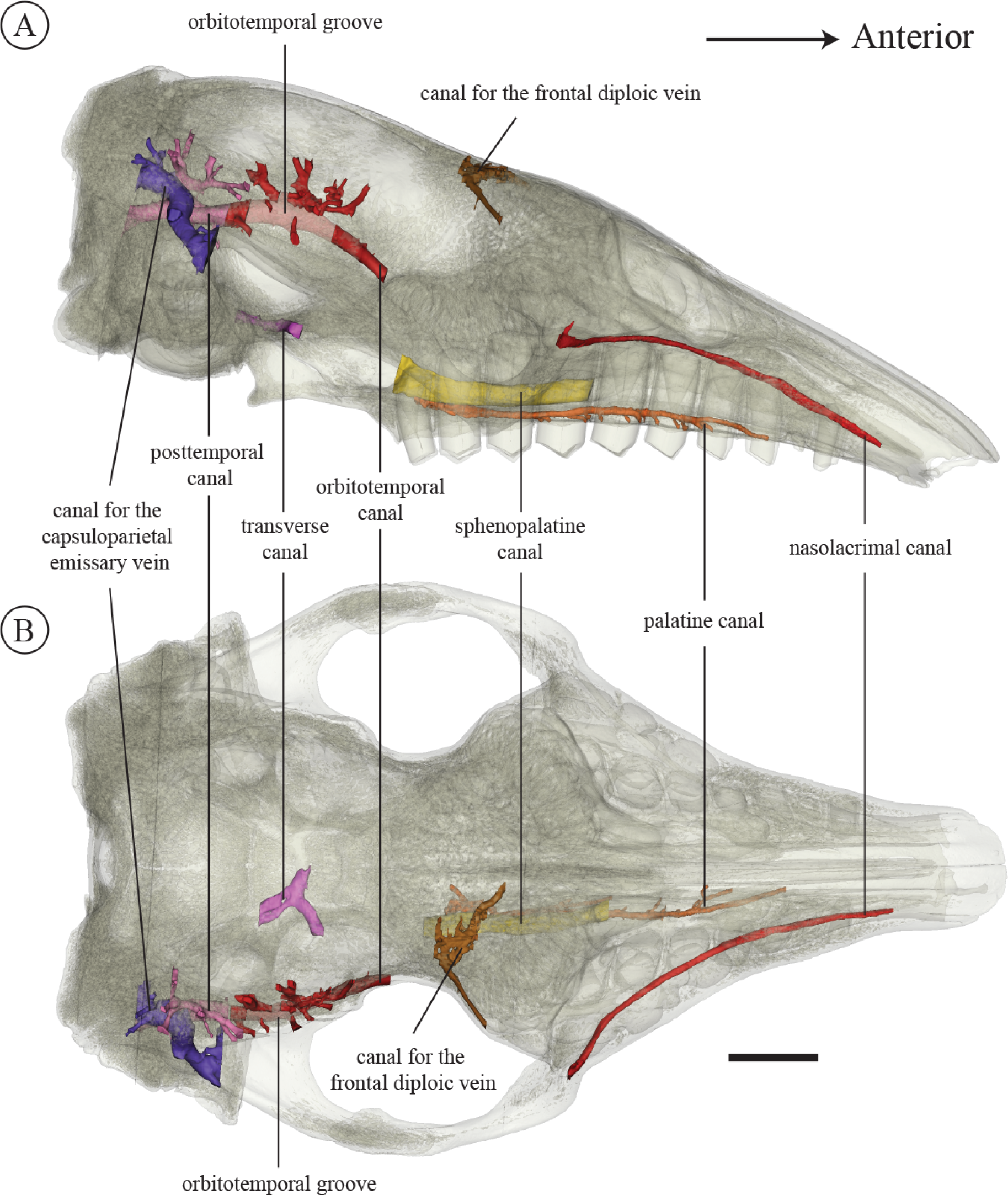
Illustration of the selected internal canals on the transparent cranium of *Euphractus sexcinctus* AMNH 133304 in right lateral (A) and ventral (B) views. Scale = 1 cm.

### 2.3 Virtual reconstruction of the selected regions

When present, the most dorsal part of the teeth did not always fill the whole alveolar cavity, leaving a void between the roof of the alveolar cavity and the dorsal edge of the tooth. Because we selected the internal orientation and curvature of the whole dental row for study, the reconstruction of alveoli was preferred over the modelling of teeth, which were sometimes absent for our specimens. In some cases, the distinction between the limit of the alveolar wall, the tooth and the sediment were not identifiable. In this case we modelled only the distinguishable structures to allow comparison (e.g., *Doellotatus* in Figure 2).

**FIGURE 2.**
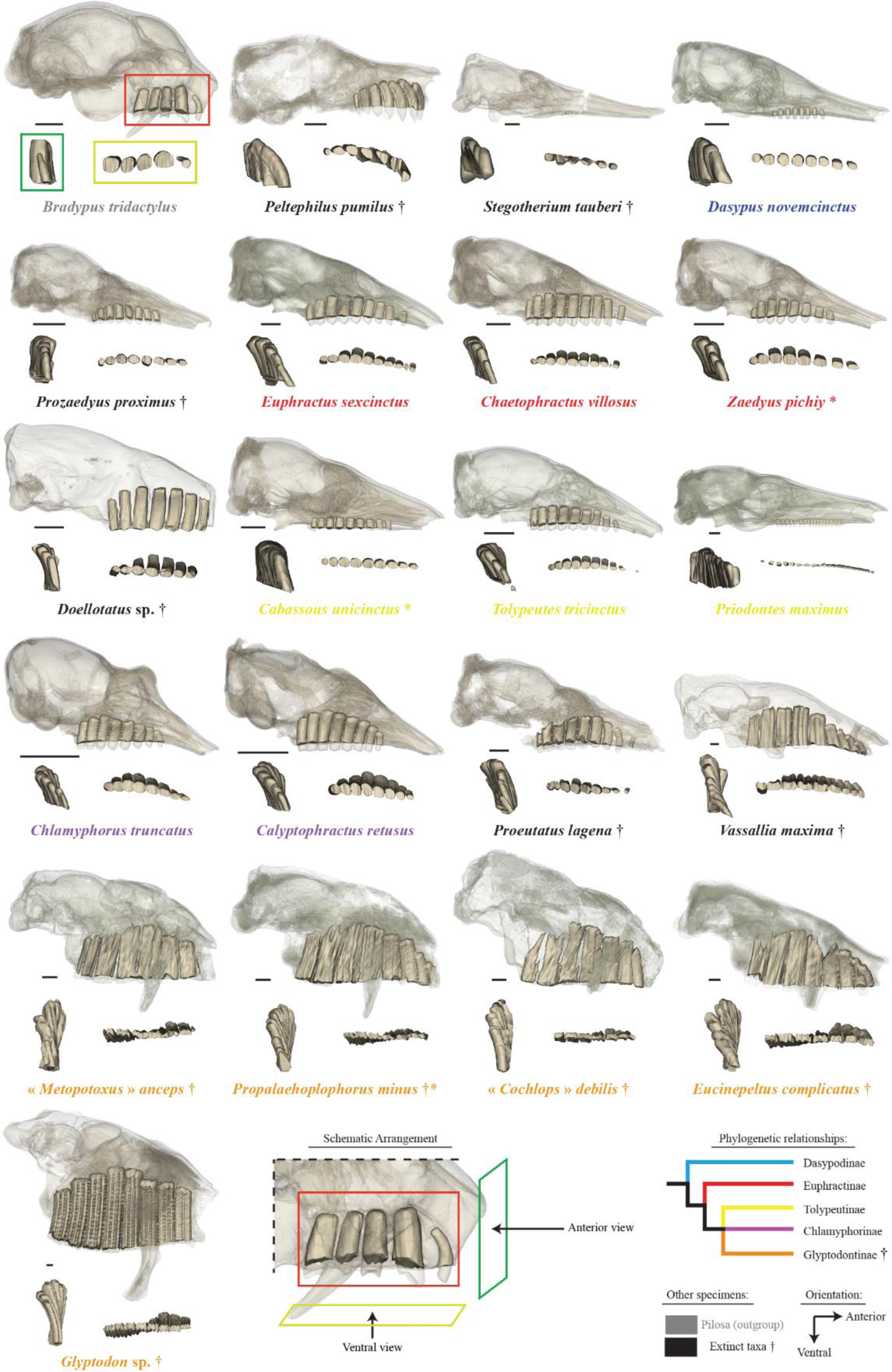
Diversity of the dental alveolar cavities in our cingulate sample in lateral (red square), ventral (yellow square) and anterior (green square) view. Crania are reconstructed with transparency to leave the cavities apparent. Colors of species names refer to their attribution at the subfamilial level according to molecular analyses (lower right panel); extinct species not allocated to one of these subfamilies are shown in black. † represents an extinct taxon. Scale = 1 cm.

Within the cranium, a canal corresponds to a duct completely enclosed in one or more bones, and which usually provides passageway to vessels and/or nerves. In some cases, a canal may not be continuously enclosed by bone, and thus turns into a groove for part of its course (e.g., orbitotemporal canal). In such a case, we have reconstructed the course of the groove in a manner similarly to that of a canal, but we illustrated the areas corresponding to the groove with increased transparency. In some cases, two canals were confluent in such a way that it was no longer possible to distinguish which part of the duct corresponds to the passage of which vessel or nerve. These regions of confluence were specified in each of our reconstructions by a shaded area (e.g., posttemporal canal and canal for the capsuloparietal emissary vein). Most selected canals were continuous with external or internal foramina. Figure S1 shows the location of each of these foramina in *Euphractus*.

### 2.4 Measurements

A few specific measurements were taken for comparison where quantification seemed to offer a better account of the observed variation than a qualitative description. Cranium length, height and width were measured to calculate the geometric mean of each specimen (= geometric mean, an estimator of the size of a specimen defined by the cubic root of the product of the three variables (Claude, 2008)). The geometric mean was log transformed to facilitate graphing of the data (Claude, 2008). Several other cranial measurements were selected, and their variation was compared to that of the geometric mean in our sample (Table S4 and Figure S2). These measurements were selected in order to quantify the following aspects that appeared interesting to us after our anatomical observations: i) the relative height of the dental row (i.e., GDRH/GCH = the ratio between the greatest dental row height and the greatest cranium height); ii) the angle between the straight line marked by the ventral most point of the tentorial process and the dorsal most point of the annular ridge (see Wible & Spaulding, 2013) and the horizontal anteroposterior axis defined here by the straight line between the mesial edge of the first tooth and the distal edge of the last tooth in sagittal view. This served to compare the internal vault inclination (IVI angle) among taxa. These measurements are further explained and summarized in Figure S2; they were taken using the linear distance and angle tools of MIMICS v. 21.0 software.

### 2.5 Reconstructing evolutionary scenarios for intracranial traits

In order to visualize possible evolutionary scenarios for the discrete and continuous intracranial traits defined based on our observations, we reconstructed a cladogram focused on the taxa in our sample by performing a parsimony analysis using PAUP v. 4.0a167 (Swofford, 2002). We used the morphological matrix of Billet et al. (2011) (largely based on that of Gaudin & Wible, 2006), which we limited to the genera in our sample. The initial matrix included all the genera of our sample except for one extant genus (*Calyptophractus*), along with four fossil genera of glyptodonts: “*Cochlops*”, *Eucinepeltus*, “*Metopotoxus*”, and *Glyptodon*. We have retained the quotation marks for the genera “*Cochlops*” and “*Metopotoxus*” in agreement with the notes of Scott (1903). They have otherwise not been revised since Scott’s (1903) report. We scored these taxa (i.e., *Calyptophractus* and several glyptodonts) based on our CT-scanned specimens and added them to the analysis. After this stage, the matrix included 22 taxa scored for 125 characters (Supporting information 1). We performed the same analysis as Billet et al. (2011), with a heuristic search involving 1000 random addition replicates, with 27 morphological characters treated as ordered and treating taxa with multiple states as polymorphic (Supporting information 1). Following the recent molecular works of Delsuc et al. (2016) and Mitchell et al. (2016), we enforced a backbone constraint similar to that of Mitchell et al. (2016). Because this resulted in an unusual position for *Proeutatus* (i.e., as the sister group of euphractines), contrary to its usual placement close to the glyptodonts in morphological analyses (Engelmann, 1985; Gaudin & Wible, 2006; Billet et al., 2011; Herrera et al., 2017), we added a constraint on this taxon to assign it a priori to a position close to glyptodonts (Figure S3A).

By using this approach, our aim was only to obtain a more consensual topology, based on recent morphological and molecular analyses, in order to discuss the relevance of our characters. This study is thus not intended to produce a new phylogenetic analysis. The strict consensus of the most parsimonious trees (with two polytomies – extant euphractines and glyptodonts except for “*Metopotoxus*”) obtained was then used to calculate branch lengths in order to explore evolutionary scenarios for our qualitative and quantitative observations over the entire topology of the tree. The strict consensus was used as a baseline cladogram (Figure S3B – the strict consensus of the same analysis without constraint is illustrated in Figure S3C for indication). As the optimization options of the analysis can affect the length of the branches, we have chosen to favor the hypothesis of convergence (= DELTRAN) rather than reversion because they have been regarded as more likely (Wake et al., 2011). For the reconstruction of evolutionary scenarios for intracranial discrete or continuous traits, one needs to complete missing data (n = 5 on 8 characters) and resolve polytomies. For the latter, we used the *multi2di* function of the *ape* package of R (Paradis & Schliep, 2019). The function resolves polytomies by adding one or several additional node(s) and corresponding branch(es) of length 0. The duplicate nodes thus have the same values, allowing the removal of duplicates *a posteriori*. In order to facilitate the visualization of the evolutionary scenarios on the tree, we have rendered the tree ultrametric using the *force.ultrametric* function of the *phytools* package of R (Revell, 2012). For the discrete traits, the missing data (i.e., 1 taxon for characters 1 and 2 (not applicable), 7 taxa for character 4; 1 taxon for character 5 – see results) were completed according to their ancestral node optimization. In case of optimization uncertainties, we also favored convergences for the same reason mentioned previously. The reconstruction and mapping of ancestral states was performed using stochastic mapping with a symmetric condition matrix (i.e., one for which all transformation rates are considered equivalent; see Bollback, 2006; Revell, 2012). This analysis was performed using the *make.simmap* function of the *phytools* package of R, with which we produced 100 simulations for each character. For continuous traits, missing data (i.e., 2 taxa for the character 8) were estimated using an approach combining the use of multiple imputations with procrustean superimposition of principal component analysis results performed on all measurement (Table S4; Clavel et al., 2014) with the *estim* function of the *mvMORPH* package of R (Clavel et al., 2015). Then, the reconstruction of the ancestral states was performed using maximum likelihood using the *contMap* function of the *phytools* package of R (Revell, 2012).

### 2.6 Institutional Abbreviations

**AMNH**, American Museum of Natural History, New York, USA; **FMNH**, Field Museum of Natural History, Chicago, USA; **MHNG**, Muséum d’Histoire Naturelle de Genève, Genève, Switzerland; **MNHN**, Muséum National d’Histoire Naturelle, Zoologie et Anatomie comparée collections (**ZM**), Mammifères et Oiseaux collections (**MO**), fossil mammal collections, Pampean (**F.PAM),** Paris, France; **NBC**, Naturalis Biodiversity Center, Leiden, Holland; **NHM**, Natural History Museum, London,; **NMNH**, National Museum of Natural History, Smithsonian Institution, Washington, DC, USA; **UF**, University of Florida, Gainesville, USA; **YPM-VPPU**, Princeton University collection housed at Peabody Museum, Yale University, USA; **ZMB**, Museum für Naturkunde, Berlin, Germany.

## 3. RESULTS – ANATOMICAL DESCRIPTION AND COMPARISON

### 3.1 Teeth and Alveolar Cavities – Orientation, Curvature and Height

The three species sampled intraspecifically (*Dasypus novemcinctus*, *Zaedyus pichiy*, and *Cabassous unicinctus* – Table 1) do not show any ontogenetic variation in the orientation and curvature of the teeth compared to the pattern observed in adults (see below). In *Dasypus* and *Zaedyus* the relative height of teeth shows an increase from the youngest to the oldest specimens, suggesting an increase in height during ontogeny (Figure 3). In *Cabassous*, which possess a reduced dentition as *Dasypus*, no clear ontogenetic trend has been observed but we lack a stage as young as *Dasypus* and *Zaedyus* (Figure 3).

**FIGURE 3.**
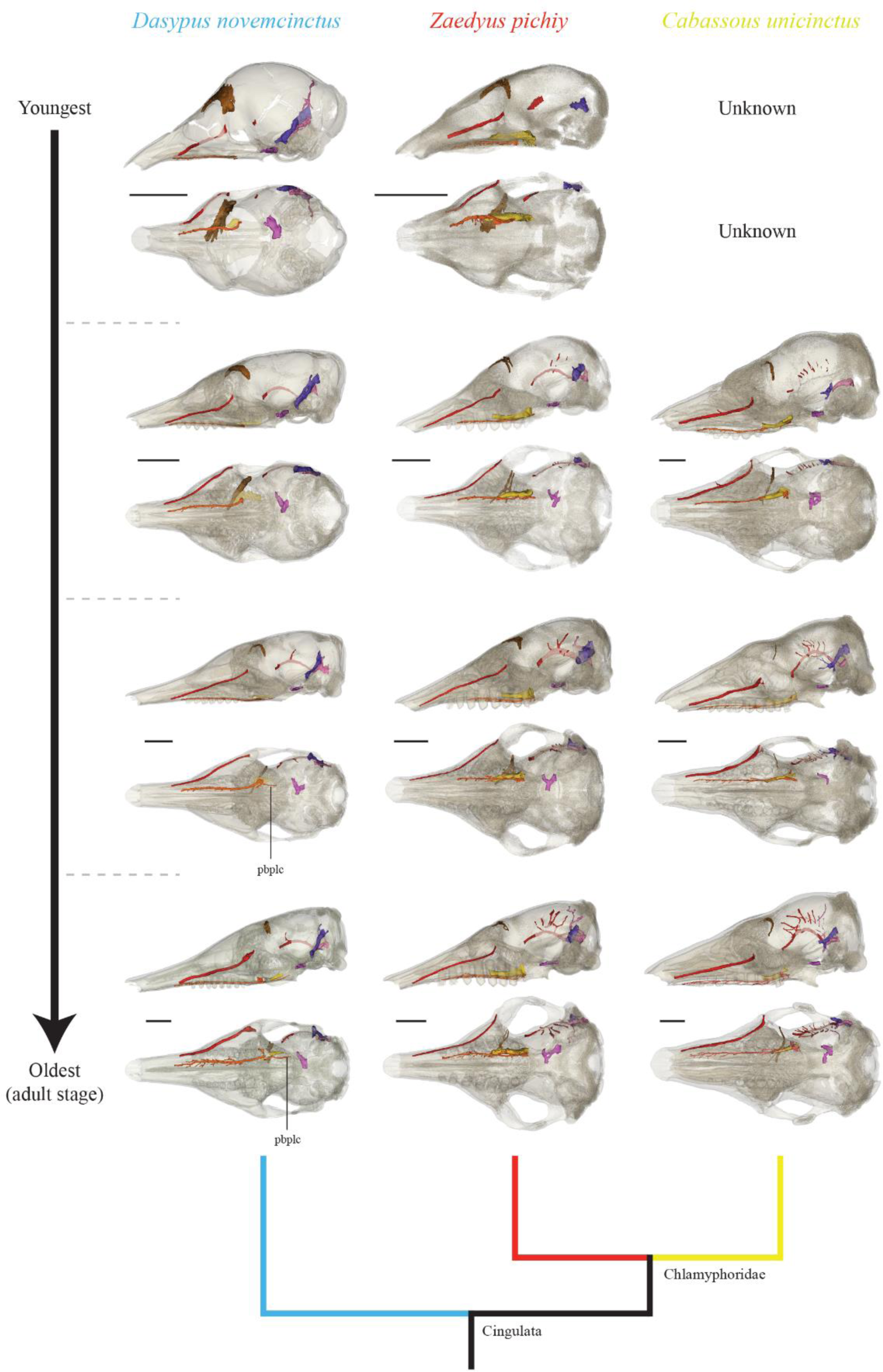
Ontogenetic variation of internal canals selected in this study viewed on transparent crania in each developmental series (*Dasypus*, *Zaedyus* and *Cabassous*). Note that early postnatal stages for *Cabassous* are not documented in our sample. See Le Verger et al. (2020) for the determination of stages. Phylogenetic relationships are indicated with colors following Figure S3. The color of canals follows that of Figure 1. Abbreviations: pbplc, posterior branch of the palatine canal. Scale = 1 cm.

*Height*--The relative height of the alveolar cavities distinguishes specimens with reduced teeth such as *Stegotherium*, *Dasypus*, *Cabassous* and *Priodontes*, from other cingulate species, some of which may even show extremely enlarged teeth, as is the case in glyptodonts (Figure 2, Table S4).

In taxa with reduced teeth (unfunctional supernumerary teeth are not considered; see González Ruiz & MacPhee, 2014; González Ruiz et al., 2014, 2015, 2017), the height of the alveolar cavity in lateral view is practically the same for all teeth (Figure 2). In the other taxa, the dorsal profile of the dental row in lateral view is curved (dorsal convexity): teeth gradually increase in height backward until the middle of the dental series and then gradually decrease in height posteriorly. The most dorsal point is reached in the middle of the dental row (at the level of Mf5 in dental rows containing 7-10 Mf) in *Euphractus* (at Mf5/Mf9), *Chaetophractus* (at Mf5/Mf9), *Doellotatus* (at Mf5/Mf9-10) and *Tolypeutes* (at 5Mf/Mf9) and more anteriorly in *Peltephilus* (at Mf2/Mf7), and *Proeutatus* (at 4Mf/Mf9). In *Bradypus*, this dorsal point is attained more anteriorly (at Mf2/Mf5). It is reached more posteriorly in *Zaedyus* (at Mf5/Mf8), *Prozaedyus* (at Mf5/Mf8), (at Mf6/Mf9), chlamyphorines (at Mf6/Mf8) and most glyptodonts (at Mf5-6/Mf8). In addition, the GDRH/GCH ratio, which express the relative dental height according to the cranium height, seems to be correlated to the body size (*R^2^* = 0.5363; *P-value* = 9.42E-06), showing that large-sized taxa have relatively taller dentitions. This is reminiscent of the increase in height of the dental row during ontogeny in *Dasypus* and *Zaedyus*. Most large-bodied taxa in the sample, i.e., *Vassallia* and glyptodonts, but not *Priodontes*, do indeed show a strongly elevated dentition in relation to the cranium depth (Figure 4). This is also the case in *Peltephilus*, for which the height of the cranium is particularly low in relation to its length. Although they are small (i.e., GCL< 43mm), chlamyphorines show a tooth height comparable in proportion to the larger-sized *Euphractus*, *Chaetophractus* and *Proeutatus* (Figure 4), meaning that they slightly depart from the detected allometric trend in having higher teeth than expected for their size. Relative tooth height is lower in *Prozaedyus*, *Zaedyus*, *Tolypeutes* and *Cabassous*, and extremely low in specimens with dental reduction – *Stegotherium*, *Dasypus* and *Priodontes* (Figure 4).

**FIGURE 4.**
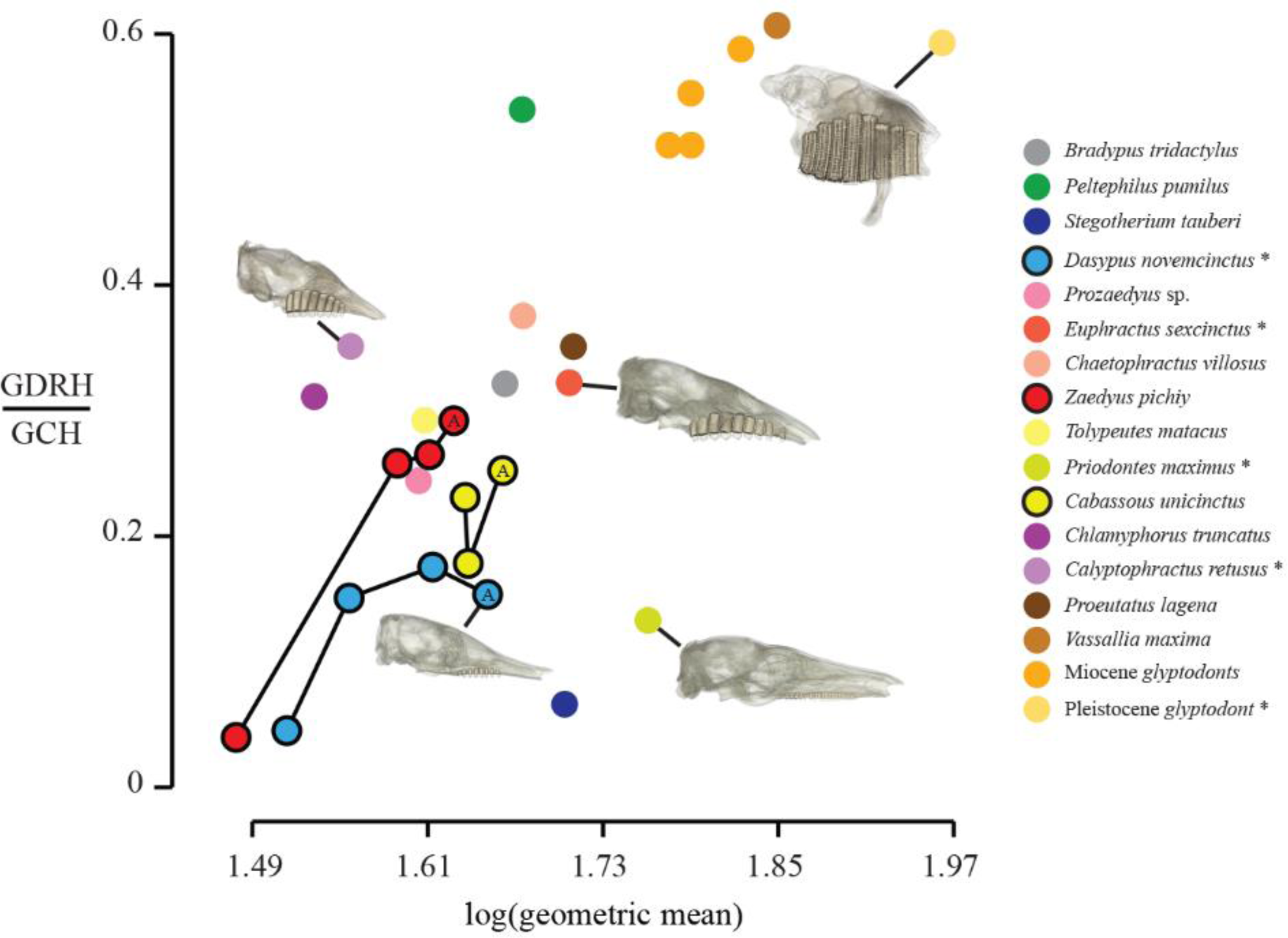
Distribution of the ratio of greatest dental row height (GDRH) to greatest cranium height (GCH) with respect to cranial size (= log(geometric mean); see Materials & Methods). For each of the three developmental series, A, symbolizes the adult specimen. Crania of specimens marked with an asterisk are illustrated in the graph. Scale = 1 cm.

*Curvature & Orientation*--As visible in anterior and ventral views (Figure 2), taxa with reduced teeth and *Peltephilus* exhibit a rather homogenous curvature and orientation of teeth along the dental row while most cingulates show gradual changes in these aspects anteroposteriorly. In most cingulates, anterior teeth are tilted lingually, i.e., tilted with a medially offset apex, while most posterior teeth often tilt labially. The tilt of posterior teeth is often much less pronounced than for anterior teeth. Similarly, taxa that show curved crowns generally show anterior teeth with an inward curvature (lingual convexity) which can be strongest in the middle of the dental row, while the two posteriormost teeth often show a lesser degree of inward curvature or even an outward curvature (labial convexity). A strong inward curvature of anterior teeth is observed in *Peltephilus*, *Doellotatus*, *Tolypeutes*, *Vassallia*, chlamyphorines, and glyptodonts in our sample, whereas other taxa show straighter crowns, and an outward curvature is observed in *Dasypus*. In lateral view, the first tooth (Mf1; P1 for *Dasypus*) or teeth show a mesial curvature (mesial convexity) and a mesially offset apex in *Bradypus*, *Peltephilus*, *Proeutatus*, *Vassallia*, *Propalaehoplophorus* and *Eucinepeltus*. Mf1 is also tilted with a mesially offset apex in *Prozaedyus*. Finally, the two to three posteriormost teeth are distinctly tilted with a distally offset apex in *Bradypus* and glyptodonts. Most other loci and taxa exhibit a nearly vertical orientation of teeth in lateral view.

Based on these observations, we propose scrutinizing evolutionary scenarios for the following two characters of the dentition (see Table 2).

> ***Character 1*** (discrete - unordered): Position of the most dorsal point of the dorsal convexity of the tooth row.
>
> **State (0)**: Toothrow not dorsally convex.
>
> **State (1)**: Anterior to middle of the tooth row.
>
> **State (2)**: At the middle of the tooth row.
>
> **State (3)**: Posterior to middle of the tooth row.

> ***Character 2*** (discrete - unordered): Curvature of anterior teeth in anterior view.
>
> **State (0)**: Inward curvature.
>
> **State (1)**: Straight.
>
> **State (2)**: Outward curvature.

### 3.2 Cranial Canals

#### 3.2.1 Frontal Diploic Vein canal

The course of this canal is entirely within the frontal bone. It opens externally at one or exceptionally two foramina located in the orbitotemporal fossa, slightly ventral to the most dorsal part of the orbital margin (Figure S1). Internally, the canal opens through one or more foramina located posteriorly to the annular ridge and near the midline without ever crossing it (Figure S4). This canal conveys the frontal diploic vein, an emissary of the dorsal cerebral vein/dorsal sagittal sinus or a vein issuing from the frontal diploë (Thewissen 1989; Wible & Gaudin, 2004; Evans & de Lahunta, 2012; Muizon et al., 2015).

In early ontogeny, the canal of the frontal diploic vein is initially extremely thick in youngest specimens of *Dasypus* (see also Billet et al., 2017) and *Zaedyus* and occupies a large part of the frontal bone (Figure 3). Its relative diameter is considerably reduced in older specimens of *Dasypus* and *Zaedyus* with no change in its curved trajectory in *Zaedyus* but becoming more strictly transverse in the adult stage in *Dasypus* (Figure 3). In *Cabassous*, it remains fairly similar from juvenile to adult stages in our sample (Figure 3).

In our adult sample, the canal is relatively straight, very thin, oriented posteromedially in dorsal view and without ramifications in *Bradypus* (Figure 5). In *Tamandua*, *Peltephilus*, *Stegotherium*, *Dasypus*, *Euphractus*, *Priodontes* and *Cabassous*, its overall course is transverse but with a posterior bend or convexity of varying degree (Figure 5). Ramifications and a posterior convexity are also found in *Prozaedyus*, *Chaetophractus*, *Zaedyus* and *Proeutatus*, but the overall course of their canal is more anteromedially oriented, as is also observed in *Stegotherium*, *Priodontes* and *Cabassous* (Figure 5). The same orientation is found in *Doellotatus* (incomplete), except that the course of the canal is completely straight and has no ramifications (Figure 5). *Tolypeutes*, on the other hand, shows a very peculiar course with a forked structure in dorsal view (Figure 5). The canal is merged with diploe in chlamyphorines but its presence is observable in *Calyptophractus* in which it has a straight and anteromedially oriented trajectory (Figure 5). This canal was not observed in *Vassallia* or in all the glyptodonts sampled. However, Gaudin (2004) notes the presence of this canal in glyptodonts and Gaudin & Lyon (2017) mention it in the pampathere *Holmesina* Simpson, 1930 (but see below). We suspect a taphonomic bias for some specimens (e.g., some glyptodonts) and call for a deeper study of this canal in cingulates before its variation can be scored.

**FIGURE 5.**
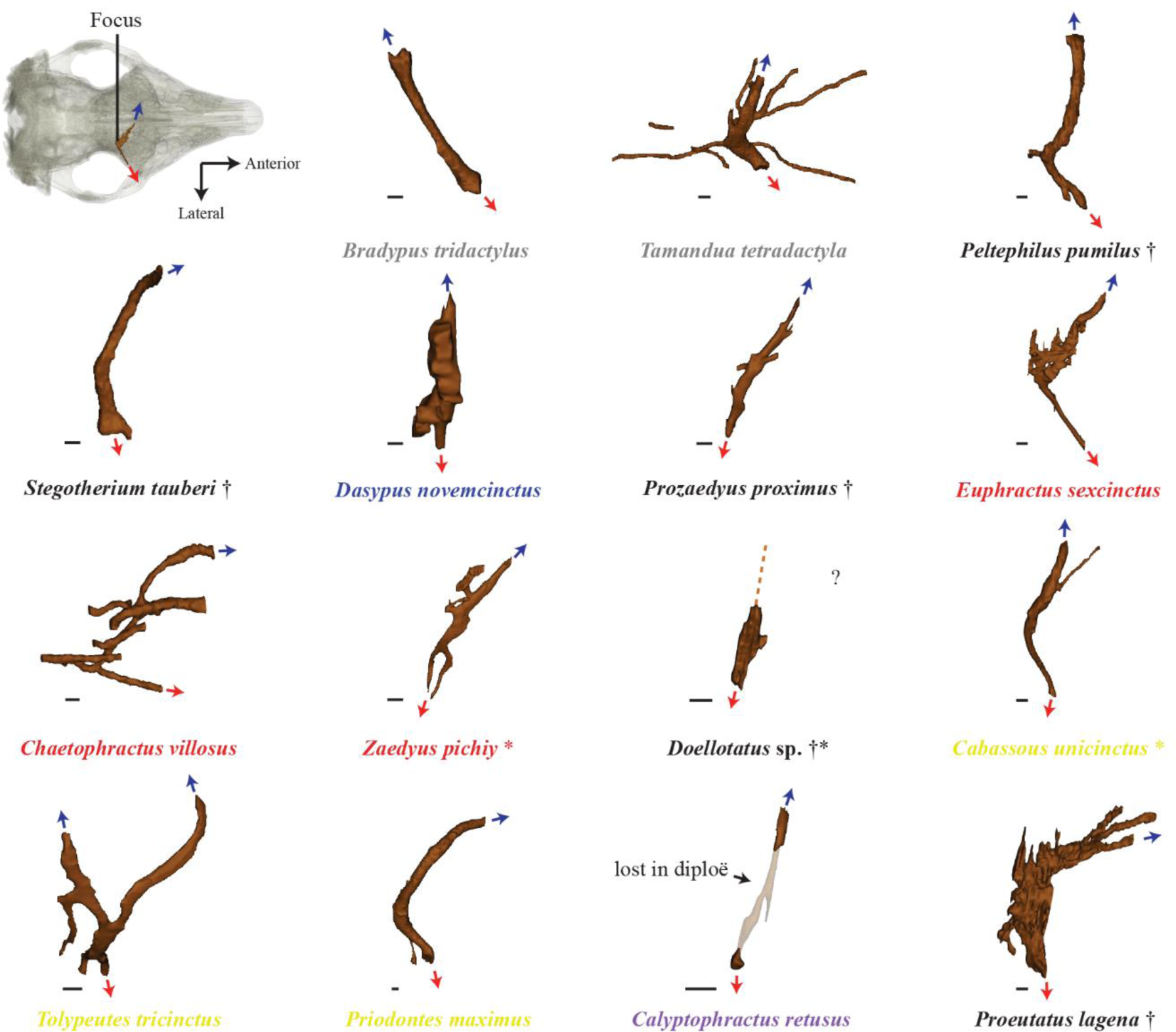
Interspecific variation of the canal for the frontal diploic vein in our sample, all illustrated in dorsal view. Red arrows mark the main external opening of the canal. Blue arrows mark the main internal path of the canal on the cranial midline. Colors of species names follow Figure 2. Scale = 1 mm.

#### 3.2.2 Nasolacrimal Canal

The nasolacrimal canal originates posteriorly at the anterior orbital edge with the lacrimal foramen (Figures 1 & S1). It runs anteriorly within the lacrimal and the maxillary bones in its posteriormost portion. More anteriorly, the canal is located between the inner wall of the maxillary and turbinates. It opens anteriorly (in front of the tooth row, except in *Peltephilus*) and ventromedially into the nasal cavity (Figures S1E & S5). This canal allows the transmission of fluids from the lacrimal sac to the nasal cavity and is potentially accompanied by a vein in *Euphractus* (see Wible & Gaudin, 2004).

The course of the nasolacrimal canal barely changes during ontogeny in our sample (Figure 3). Its length changes and follows the lengthening of the snout that accompanies the growth of the cranium (Le Verger et al., 2020). In addition, *Cabassous* and, to a lesser degree, the adult stage of *Zaedyus* show a more medial orientation of the course of the nasolacrimal canal in its most posterior part which differs slightly from *Dasypus* and other stages of *Zaedyus*, where it runs more parallel to the external surface of the cranium (Figure 3).

The position of the nasolacrimal canal and lacrimal foramen varies among taxa, although this feature was not scored in previous matrices (e.g., Gaudin & Wible, 2006). The lacrimal foramen is particularly high relative to the orbital edge (closer to the most dorsal point of the orbit than to the jugal bone in lateral view) in chlamyphorines, *Proeutatus* and glyptodonts (Figure 6). The trajectory of the nasolacrimal canal is strongly sigmoid in *Tamandua* (Figure 6). In *Bradypus*, *Peltephilus*, *Proeutatus*, *Euphractus* and *Chaetophractus*, the course of the nasolacrimal canal is slightly convex dorsally (Figure 6). The curvature is much pronounced in *Vassallia* in the posterior half of its course as it passes above the tooth row (Figure 6). In dasypodines, *Prozaedyus*, *Zaedyus*, tolypeutines, and chlamyphorines, the nasolacrimal canal is relatively straight or it bears a slight ventral convexity in its posterior part (Figure 6). In glyptodonts, the canal runs ventromedially and not anteriorly from the lacrimal foramen (Figures 6 & 7), and curves strongly inward anteriorly when approaching the sagittal plane of the cranium (Figures 6 & 7).

**FIGURE 6.**
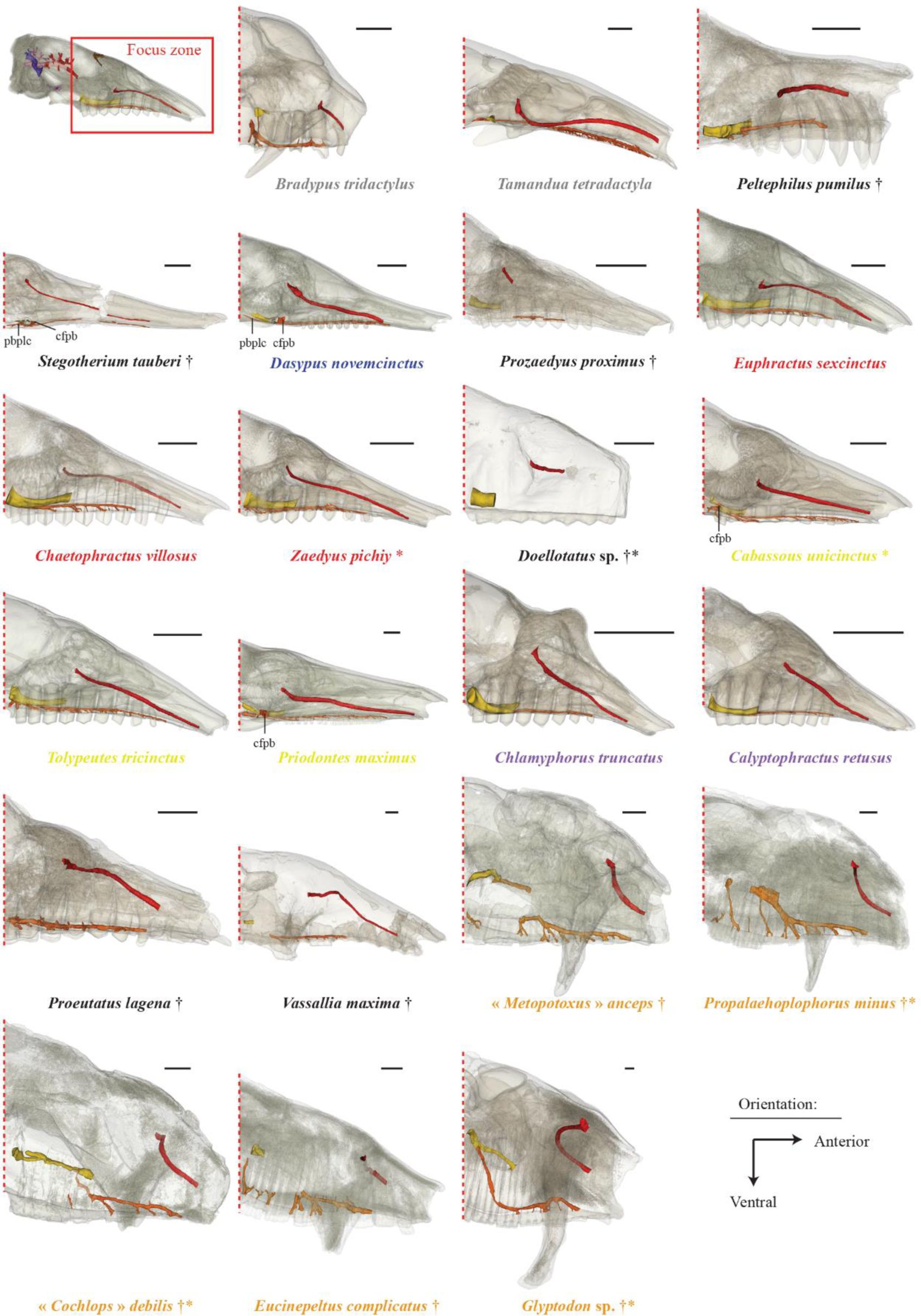
Interspecific variation in our sample of the reconstructed nasolacrimal (= red), sphenopalatine (= yellow) and palatine (= orange) canals shown in lateral view. Crania are reconstructed with transparency. Colors of species names follow Figure 2. Abbreviations: cfpb, caudal foramen for palatine canal branch; pbplc, posterior branch of palatine canal. Scale = 1 cm.

**FIGURE 7.**
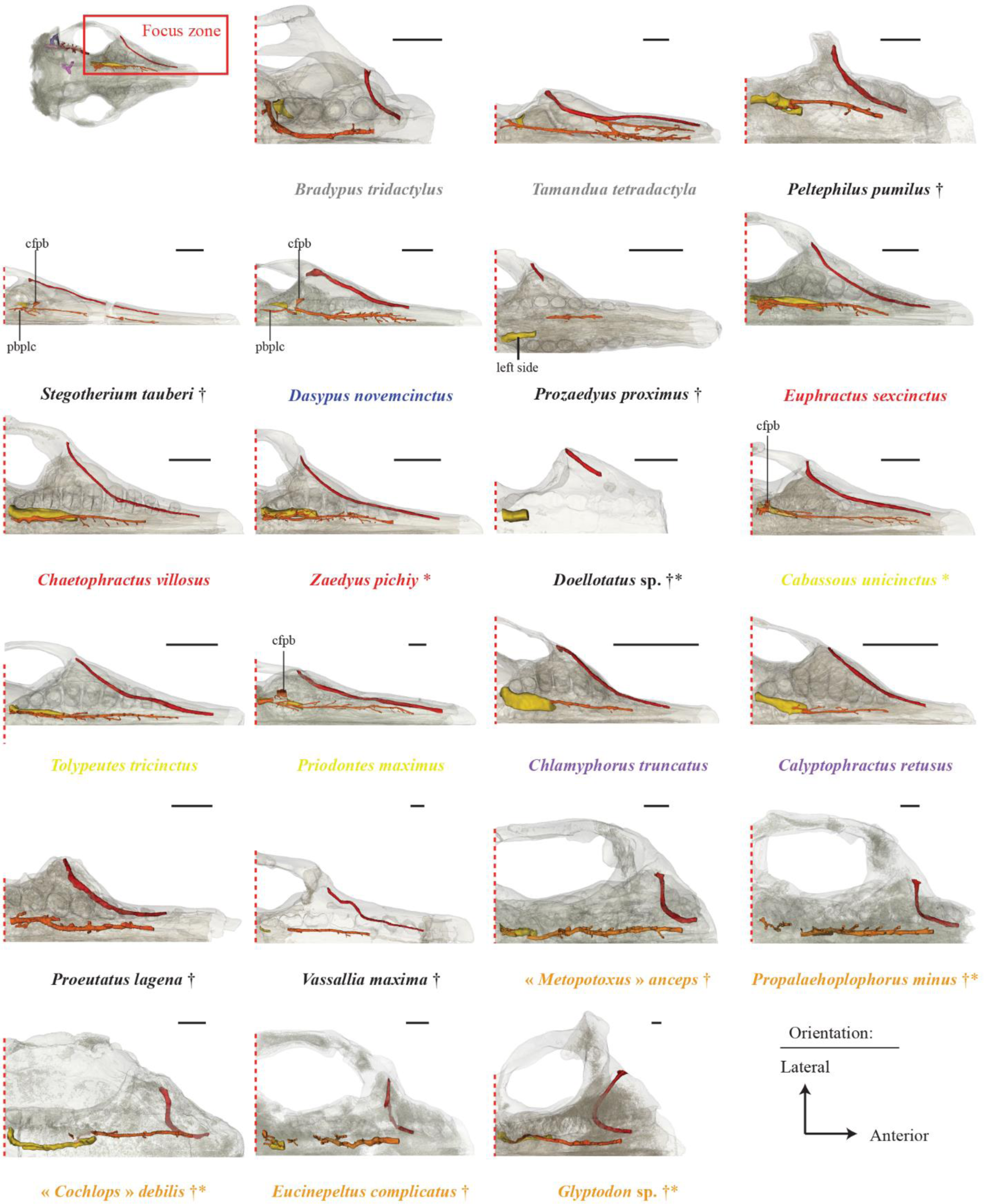
Interspecific variation in our sample of the reconstructed nasolacrimal (= red), sphenopalatine (= yellow) and palatine (= orange) canals shown in ventral view. Crania are reconstructed with transparency. Colors of species names follow Figure 2. Abbreviations: cfpb, caudal foramen for palatine canal branch; pbplc, posterior branch of palatine canal. Scale = 1 cm.

Whereas glyptodonts appear to show a unique condition in the course of their nasolacrimal canal, the anterior opening of this canal is located halfway up in their nasal cavity as in *Bradypus*, *Tamandua*, *Peltephilus*, *Doellotatus*, *Proeutatus* and *Vassallia.* In comparison, this opening is closer to the palate in other specimens (Figure 6). This variation can be scored in the following character (see Table 2).

> ***Character 3*** (discrete - unordered): Anterior opening of the nasolacrimal canal in nasal cavity.
>
> **State (0)**: Halfway up in the nasal cavity.
>
> **State (1)**: Ventral in the nasal cavity, close to the palate.

#### 3.2.3 Palatine and Sphenopalatine Canals

Two large canals are enclosed within the maxillary and palatine bones forming the hard palate in cingulates: the palatine canal and the sphenopalatine canal (Wible & Gaudin, 2004). The palatine canal is generally thinner, longer than and positioned medial to the sphenopalatine canal. The palatine canal transmits the major and minor palatine nerve, artery, and vein in *Euphractus* (Wible & Gaudin, 2004). This canal opens externally at the caudal palatine foramen posteriorly and at the multiple foramina scattered throughout the hard palate (both in the palatine and maxillary) anteriorly (Figure S1). The sphenopalatine canal, which is often very wide in cingulates, transmits the caudal nasal nerve and the sphenopalatine artery and vein (Wible & Gaudin, 2004). The sphenopalatine canal opens anteriorly in the nasal cavity and posteriorly in the pterygopalatine fossa just posterior to the dental row and close to or confluent with the caudal palatine foramen (Wible & Gaudin, 2004; Figure S1). The palatine canal and sphenopalatine canal are confluent in part of their posterior course in many of the species in our sample.

The palatine canal and sphenopalatine canal vary little during ontogeny within the three species of our ontogenetic series (Figure 3). The branch of the palatine canal posterior to its region of confluence with the sphenopalatine canal in *Dasypus* and *Cabassous* appears in the subadult (or intermediate) stage and elongates further in the adult stage (Figure 3). In the youngest specimen of *Zaedyus*, the sphenopalatine canal is much wider relative to cranium size than in later stages, and there is no region of confluence with the palatine canal (Figure 3). In the youngest *Dasypus* specimen, the sphenopalatine canal is absent. It is present only as a wide groove in the youngest to subadult stages, but is completely enclosed in the adult stage of *Dasypus* in our sample (Figure 3).

In our adult sample, the shape and trajectory of the palatine canal does not vary much among species (Figures 6 & 7). In glyptodonts, the palatine canal strongly ascends posteriorly (starting from Mf6) (Figure 6) and forms a groove running along the inner wall of the nasopharyngeal canal (Figure S6). In other cingulates, the palatine canal does not show such an ascending trajectory posteriorly. The sphenopalatine canal, which is thicker than the palatine canal, features a relatively straight trajectory in lateral view in *Peltephilus*, dasypodines, *Prozaedyus*, *Doellotatus*, *Euphractus*, *Zaedyus*, and “*Cochlops*”, whereas it shows a strong ventral curvature in *Chaetophractus*, tolypeutines, chlamyphorines and *Glyptodon* (Figure 6). It should be further noted that the dorsal position of the sphenopalatine canal relative to the nasopharyngeal canal is a feature shared by *Vassallia* and *Glyptodon* (Figure S7). In *Bradypus* and *Tamandua*, the palatine canal does not contact the sphenopalatine canal (Figures 6 & 7). The two canals are in contact in a small region of confluence in the dasypodines whereas the palatine canal becomes completely confluent with the sphenopalatine canal, and sometimes accompanied by ventrally oriented auxiliary branches of the palatine canal, in *Peltephilus*, tolypeutines, chlamyphorines and *Glyptodon* (Figures 6 & 7). In euphractines, the confluence between the sphenopalatine canal and the palatine canal is not as clear-cut as in the above-cited cingulates (Figures 6 & 7). The palatine canal of euphractines runs ventrally along the sphenopalatine canal in the same figure 8-shaped canal while being completely conjoined only in their most posterior part and at their posterior opening (Figures 6 & 7). The extent of this confluence is highly variable in *Euphractus* according to Wible & Gaudin (2004), and in other cingulates as well (Gaudin & Wible, 2006 – character 71). We observed the same variation as Gaudin & Wible (2006) concerning the extent of this confluence in the vicinity of the posterior opening of the sphenopalatine canal for all the taxa in our sample. Several branches may emerge from this region of confluence (Figures 6 & 7), some of which present an interesting pattern. One branch opens externally from the caudal foramen of the palatine canal in dasypodines, *Priodontes* and *Cabassous* (Figures 6 & 7). A second branch extends posteriorly into the hard palate, before disappearing in the bone diploe in dasypodines. In *Euphractus*, *Zaedyus*, *Priodontes*, and *Cabassous*, two or more branches extend posteriorly into the hard palate, but they open in the nasopharyngeal canal (Figures 6 & 7). Several other ramifications of the palatine canal are present in our sample, but it remains to be determined how variable these are at the intraspecific level with a dataset larger than that of Table S1.

**FIGURE 8.**
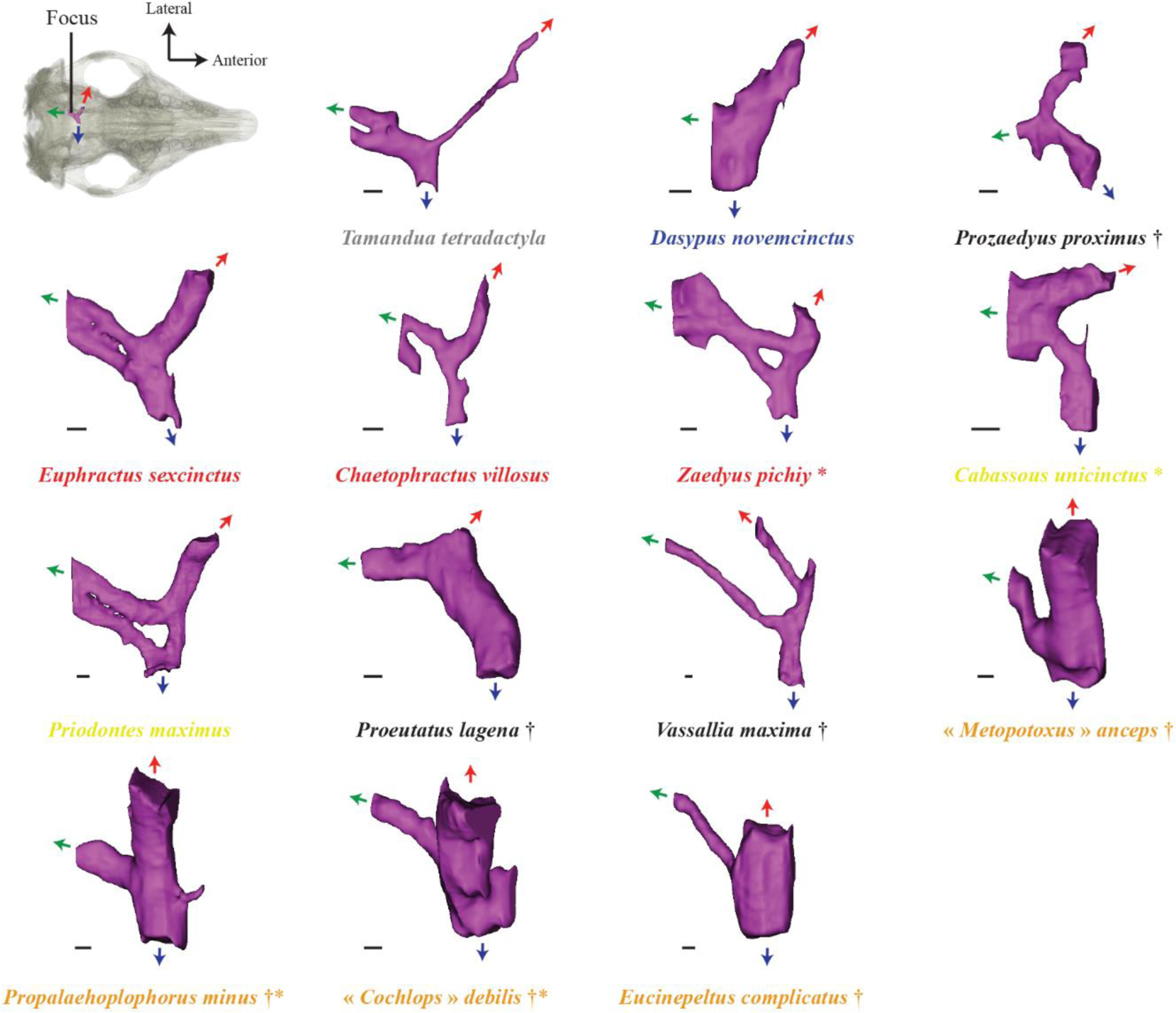
Interspecific variation of the transverse canal in ventral view in our sample. Red arrows mark the main external opening of the canal. Blue arrows mark the branch crossing the midline. Green arrows mark the main posterior branch which connects to the inferior petrosal sinus (see text). Colors of species names follow Figure 2. Scale = 1 mm.

Based on the observation of these two canals, we proposed to scrutinize evolutionary scenarios for the following character (see Table 2).

> ***Character 4*** (discrete - unordered): Sphenopalatine and palatine canal connection.
>
> **State (0)**: No contact.
>
> **State (1)**: Partial contact, with palatine canal running ventrally along the sphenopalatine canal and producing posterior branches.
>
> **State (2)**: Complete fusion.

#### 3.2.4 Transverse Canal

When present, the transverse canal is marked externally by a foramen positioned anteroventrally to or within the foramen ovale in the alisphenoid (Figure S1). This canal crosses the cranium transversally at the level of the basisphenoid and transmits a large vein issued from the cavernous sinus (Sánchez-Villagra & Wible, 2002; Wible & Gaudin, 2004). Often, one or more branches originate from this canal and extend posteriorly towards the lateral edge of the basisphenoid to join the inferior petrosal sinus (Wible & Gaudin, 2004).

The transverse canal appears early in *Dasypus* and varies little during ontogeny. Although, in *Zaedyus* and *Cabassous*, the connection between the transverse canal and the posterior branch extending towards the inferior petrosal sinus varies in both series, we have not observed a clear ontogenetic pattern of variation in our three species (Figure 3).

In *Bradypus*, *Peltephilus*, *Stegotherium*, *Doellotatus*, *Tolypeutes* and *Glyptodon*, there is no transverse canal, or it is not detectable (scored as present in *Doellotatus* and *Stegotherium*, variable in *Tolypeutes* by Gaudin & Wible, 2006 – character 111). Chlamyphorines have a transverse canal foramen opening directly within the braincase, and thus there is a very short canal that does not cross the midline of the cranium. *Tamandua* stands out in having a thin lateral part of the transverse canal which is not comparable to the condition in Cingulata (Figure 8). In *Dasypus*, *Prozaedyus*, euphractines, *Priodontes* and *Cabassous*, the branch starting from the transverse canal foramen is oriented posteromedially in ventral view (Figure 8). In the Miocene glyptodonts, it is medially oriented (Figure 8). In *Vassallia* it is anteromedially oriented (Figure 8). This lateral branch of the transverse canal splits into a branch that crosses the midline in the basisphenoid bone and a branch that joins the inferior petrosal sinus (Figure 8). In *Proeutatus*, this bifurcation occurs much closer to the transverse canal foramen than in other taxa (Figure 8). It is relatively more medial in *Prozaedyus*, euphractines, *Priodontes* and *Cabassous* (Figure 8). In *Tamandua*, *Dasypus*, *Vassallia* and Miocene glyptodonts, this bifurcation is closer to the sagittal plane than to the foramen (Figure 8). The transverse medial branch is anteromedially oriented in ventral view in *Prozaedyus*, *Cabassous* and *Proeutatus*, whereas it is medially oriented in the other taxa (Figure 8). Our specimens of *Tamandua*, euphractines (not complete in *Chaetophractus*) and *Priodontes* show an additional branch connecting the transverse medial branch to the posterior branch reaching the inferior petrosal sinus (Figure 8). In *Vassallia* and the Miocene glyptodonts, the posterior branch reaching the inferior petrosal sinus is much thinner than the rest of the canal, in contrast to the other taxa, in which the whole canal exhibits a relatively homogeneous width (Figure 8).

Based on these observations of the transverse canal, we propose to scrutinize evolutionary scenarios for the following character (see Table 2).

> ***Character 5*** (discrete - unordered): Orientation of the branch starting from the transverse canal foramen.
>
> **State (0)**: Posteromedial.
>
> **State (1)**: Medial or anteromedial.
>
> **State (2)**: No canal.

#### 3.2.5 Orbitotemporal and Posttemporal Canals

Within the braincase, the orbitotemporal canal provides passageway to the rostral extension of the *ramus superior* of the stapedial artery, or orbitotemporal artery (giving rise to the *ramus supraorbitalis* in the orbital region), and a few small veins (Wible & Gaudin, 2004) whereas the more posterior posttemporal canal transmits the *arteria diploëtica magna* and the large *vena diploëtica magna* (Wible & Gaudin, 2004). These two canals can give rise to the arterial and venous *rami temporales* along the lateral wall of the braincase which exit via numerous foramina on the cranial roof to irrigate the *temporalis* muscle (Wible & Gaudin, 2004; Figure S1). The posttemporal canal is connected posteriorly to an external groove in the petro-occipital region marking the passage of the occipital artery (Wible & Gaudin, 2004; Figure S1). From there, the posttemporal canal extends forward as a canal enclosed between the petrosal and squamosal (or only in the squamosal) and oriented mostly horizontally. More anteriorly, past the petrosal, the posttemporal canal is connected to the orbitotemporal canal, which runs further anteriorly within or on the inner surface of the lateral braincase wall (formed by the squamosal, frontal and sometimes the parietal). The delimitation between the two canals generally occurs where the canal for the *ramus superior* joins them (Muizon et al., 2015). However, we were only able to observe a canal possibly transmitting the *ramus superior* in *Tolypeutes* (Figure S8), but not in the other specimens. Consequently, we followed the suggestion of Wible & Gaudin (2004), who separate the two canals at the level of the postglenoid foramen. In some taxa, the orbitotemporal canal is not completely enclosed for parts of its length, and appears instead as a groove on the internal wall of the braincase. The anterior opening of the orbitotemporal canal is located in the orbitotemporal region, just posteroventral to the postorbital constriction (Figure S1; Wible & Gaudin, 2004; = cranioorbital foramen in Gaudin & Wible, 2006; see also Muizon et al. (2015) for multiple illustrations of these canals in another placental taxon).

The orbitotemporal canal only appears as a short canal near the orbitotemporal region in the youngest specimens of all three species, whereas the canal (or groove) is clearly visible for its entire length in later stages (Figure 3). The connections of the orbitotemporal canal with canals for the numerous *rami temporales* occur in the two last stages in both *Zaedyus* and *Cabassous* (Figure 3). The posttemporal canal is already well formed in the youngest specimen of *Dasypus* and hardly changes during ontogeny (Figure 3). In *Zaedyus*, the posttemporal canal is not enclosed in the youngest stage. In *Cabassous*, the posttemporal canal is present in the youngest stage sampled, but it expands anteriorly during the two subsequent stages (Figure 3).

In our adult sample, *Bradypus*, *Tamandua*, euphractines, *Vassallia* and glyptodonts possess a posttemporal canal emerging at the center of the occiput in lateral view and at the level of the jugular foramen (Figure 9). In *Peltephilus*, it originates more ventrally, even with the most dorsal margin of the occipital condyles in lateral view (Figure 9). In dasypodines, tolypeutines and chlamyphorines, the posterior extremity of the canal is at an intermediate height (Figure 9). Apart from the position of the posterior opening of this canal (= posttemporal foramen of Wible & Gaudin, 2004), we were unable to determine any systematically significant variation regarding its direction, length, or width.

**FIGURE 9.**
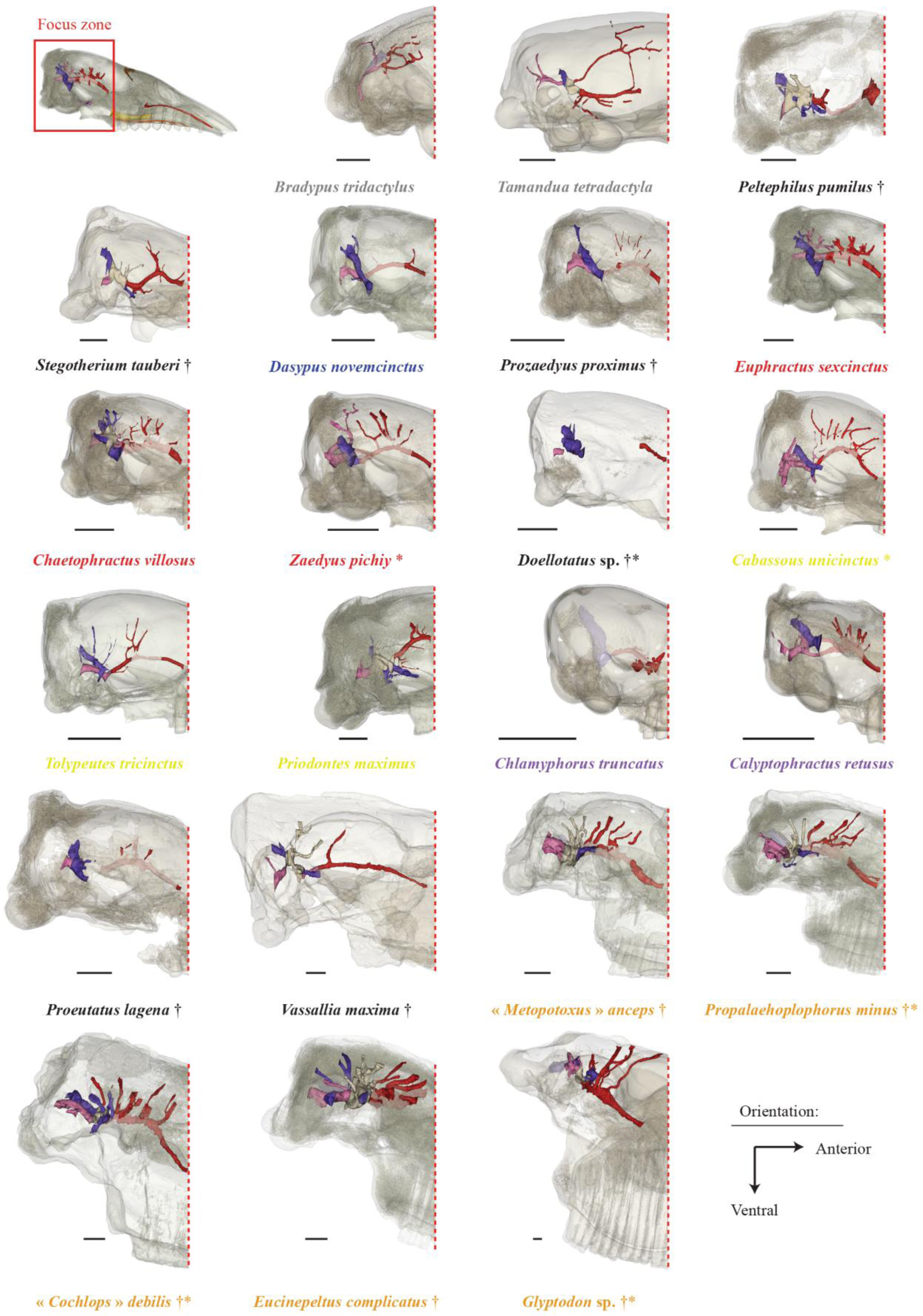
Interspecific variation in our sample of the orbitotemporal (= red), posttemporal (= pink) and capsuloparietal emissary vein (= blue) canals shown in lateral view. Caudal part of the cranium is transparent. Parts of the canals showing transparency symbolize a groove instead of a fully enclosed canal. Colors of species names follow Figure 2. Scale = 1 cm.

**FIGURE 10.**
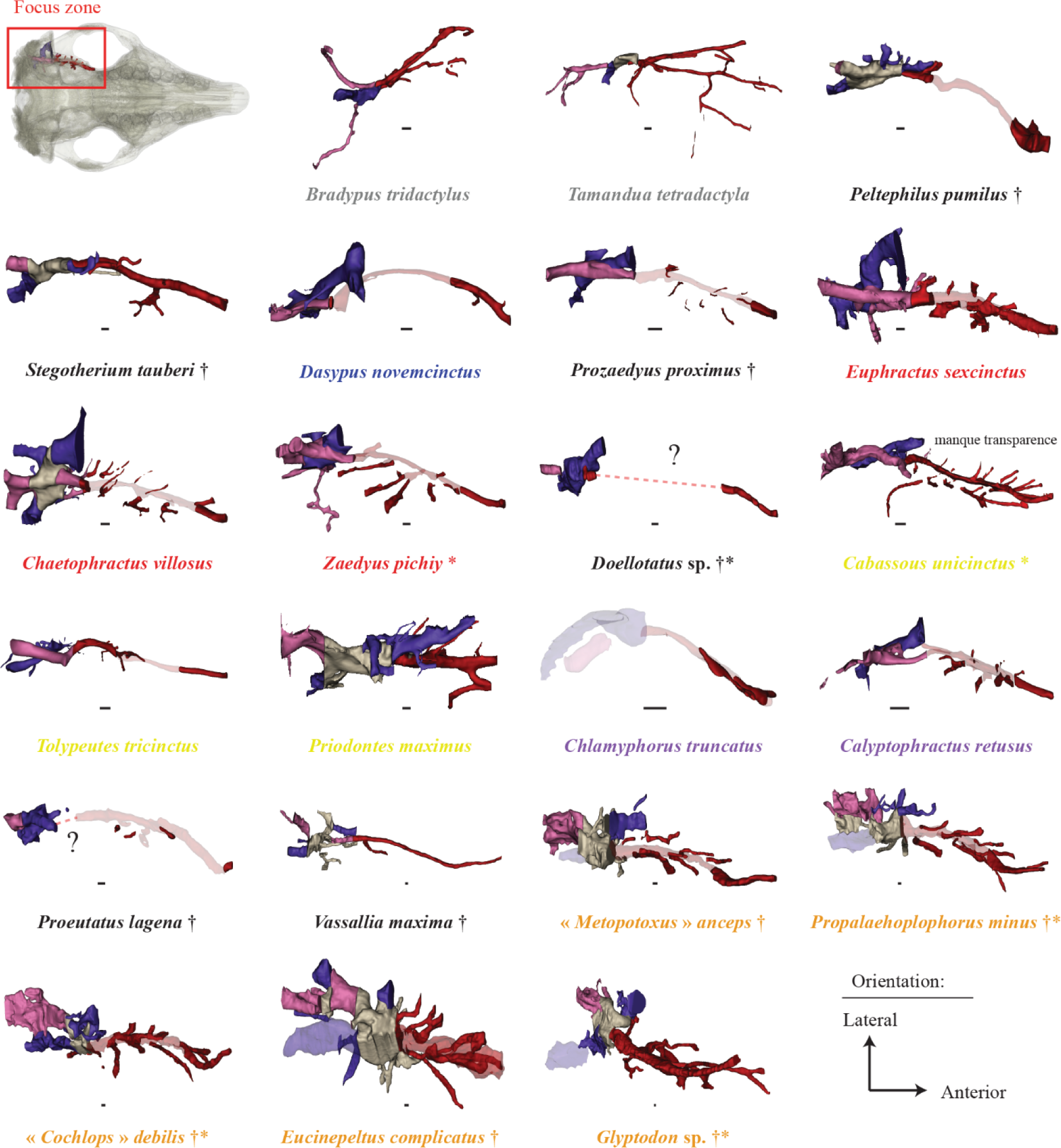
Interspecific variation in our sample of the orbitotemporal (= red), posttemporal (= pink) and capsuloparietal emissary vein (= blue) canals shown in ventral view. Transparent parts of the canals indicate a groove instead of a fully enclosed canal. Colors of species names follow Figure 2. Scale = 1 mm.

In *Bradypus* and *Tamandua*, the orbitotemporal canal does not reach the orbitotemporal region (Figures 9 & 10). It reaches the orbitotemporal region in all the cingulates in our sample except for *Priodontes* (Figures 9 & 10). Gaudin & Wible (2006) mention the absence of the anterior opening of the orbitotemporal canal in *Priodontes* (character 78), but also in *Chlamyphorus* and *Vassallia* for which we observe a clear opening in the orbitotemporal region (Figure 9). In *Bradypus*, *Tamandua* and *Peltephilus*, this canal bears a ventral curvature (strong in *Peltephilus*), whereas its trajectory exhibits a more or less strong dorsal curvature in all the other taxa (Figure 9). In *Peltephilus*, *Dasypus*, *Prozaedyus*, euphractines, *Tolypeutes*, *Cabassous*, chlamyphorines, *Proeutatus* and Miocene glyptodonts, the orbitotemporal canal becomes a groove on the internal lateral wall of the braincase before being enclosed again as a canal close to its external anterior opening (Figure 9). In *Calyptophractus*, *Proeutatus* and glyptodonts, the anterior half of the orbitotemporal canal displays a pronounced downward trajectory, and its anterior opening is located lower than its posterior connection with the posttemporal canal (Figure 9). *Calyptophractus* stands out, however, in having two branches that connect the orbitotemporal and posttemporal canals (Figure 9). A large region of confluence between the orbitotemporal and posttemporal canals and the canal for the capsuloparietal emissary vein is identified in *Tamandua*, *Peltephilus*, *Stegotherium*, *Dasypus*, *Chaetophractus*, *Priodontes*, *Vassallia* and the glyptodonts (and potentially also in *Proeutatus* but the orbitotemporal canal is not distinct in this region; Figures 9 & 10). This confluent region is reminiscent of the petrosquamosal fossa in *Alcidedorbignya* Muizon & Marshall, 1987 (Muizon et al., 2015). The orbitotemporal and posttemporal canals connect with the *rami temporales* through multiple small canals whose number remains quite variable at the intraspecific level. However, a high density of *rami temporales* for the orbitotemporal canal is particularly noticeable in euphractines, *Cabassous*, *Calyptophractus* and glyptodonts (Figure 9). This density is coded by Gaudin & Wible (2006), who counted the number of foramina for *rami temporales* in the temporal fossa of parietal (0, equal to or less than five; 1, greater than five – character 97).

In their matrix, almost all cingulates are coded as having more than 5 foramina. The variations in the density of the canals for the *rami temporales* observed in our study call for further investigation of these structures, which could potentially carry an interesting phylogenetic signal among cingulates.

Based on these observations of the orbitotemporal canal, we propose to scrutinize evolutionary scenarios for the following character (see Table 2).

> ***Character 6*** (discrete - unordered): Downward trajectory of the orbitotemporal canal.
>
> **State (0)**: Canal does not reach the orbitotemporal region.
>
> **State (1)**: The anterior part of the canal has a slight downward trajectory ending at the same or almost the same height as its origin from the posttemporal canal.
>
> **State (2)**: The canal has a strong downward anterior trajectory, and its anterior opening is much lower than its posterior connection to the posttemporal canal.

#### 3.2.6 Canal of the Capsuloparietal Emissary Vein

The canal for the capsuloparietal emissary vein opens anteroventrally at the postglenoid and suprameatal foramina, and connects to the groove for the transverse sinus posterodorsally (Wible, 1993; Muizon et al., 2015). From the cranial roof, the transverse sinus runs lateroventrally in a wide groove excavated in the inner surface of the parietal and extends anteroventrally within a canal formed by the petrosal medially and the squamosal laterally. This canal opens externally through the suprameatal foramen, the postglenoid foramen and accessory foramina at the posterior base of the zygomatic arch (e.g., Wible & Gaudin, 2004; Figure S1). It transmits the capsuloparietal emissary vein (= retroarticular vein in *Canis* Linnaeus 1758 (Evans & de Lahunta, 2012); postglenoid vein in *Cynocephalus* Boddaert 1768 (Wible, 1993) and *Dasypus* Linnaeus 1758 (Wible, 2010)). In our sample, this canal is often partly confluent with the posttemporal and orbitotemporal canal in its posterodorsal portion (Figure S9).

The canal for the capsuloparietal emissary vein is already well formed in the youngest specimen of *Dasypus* and hardly changes during ontogeny, except that it becomes relatively thinner (Figure 3). In *Zaedyus*, it is only partly formed in the youngest specimen, and its trajectory is better marked in older stages (Figure 3). In the youngest *Cabassous*, this canal is noticeably short and only marked in its most ventral part (glenoid region). It is only in older stages that the canal elongates posterodorsally (Figure 3).

The canal is absent and the passage of the vein is only indicated by a groove in *Bradypus* and *Chlamyphorus*. The groove is very wide in the latter two (Figure 9). The canal of the capsuloparietal emissary vein opens through the postglenoid foramen and potentially in the suprameatal foramen as well (see *rami temporales* opening in the squamosal whose number varies in our intraspecific sampling (Table S1)) in *Peltephilus*, *Prozaedyus*, *Doellotatus*, euphractines, chlamyphorines, *Proeutatus*, *Vassallia* and the glyptodonts (as notably scored for the glenoid region by Gaudin & Wible, 2006 – character 119; Figures 9 & 10). In tolypeutines, the canal always opens by a suprameatal foramen with a well-marked branch, whereas its opening via the postglenoid foramen may sometimes be absent, as we have occasionally observed the absence of a postglenoid foramen in our intraspecific sample of *Cabassous* (Table S1). For all our taxa except *Bradypus*, the canal extends anteroventrally the cranial roof (Figure 9). However, its inclination varies considerably from one species to another. It is steeply inclined in dasypodines, slightly less so in *Prozaedyus*, *Doellotatus*, euphractines and *Proeutatus*, and much less so in tolypeutines, *Calyptophractus*, *Vassallia* and glyptodonts (Figure 9). The canal is sinuous in *Euphractus* and *Chaetophractus*. In glyptodonts, the canal even has an almost anteroposterior orientation near its anterior opening (Figures 9 & 10). The canal (or groove) for the capsuloparietal emissary vein reaches a region of confluence (= petrosquamosal fossa, Muizon et al., 2015) with the other canals of the braincase in *Tamandua*, *Peltephilus*, dasypodines (particularly thin in *Dasypus*), *Chaetophractus*, *Priodontes*, *Vassallia* and glyptodonts (Figures 9 & 10). For the other taxa, the canal for the capsuloparietal emissary vein is only adjacent to the other canals (Figures 9 & 10 – see Figure S9 for an illustration of the different cases in our sample). The variation in length and thickness of this canal does not provide clear systematic information. A relatively short canal is present in *Tamandua*, *Doellotatus*, *Zaedyus*, and *Proeutatus* compared with dasypodines (Figure 9). Chlamyphorines and *Zaedyus* show a relatively thicker canal, as compared with the very thin canal in *Vassallia*. The fact that small taxa have a relatively thicker canal is reminiscent of young specimens of *Dasypus* as compared to older and larger specimens of the same species (Figure 3) and suggest a potential allometric pattern.

Based on these observations, we propose to scrutinize evolutionary scenarios for the following character related to the anterolateral extremity of the canal (see Table 2).

> ***Character 7*** (discrete - unordered): Canal of the capsuloparietal emissary vein inclination in lateral view.
>
> **State (0)**: Anterodorsal orientation or canal very short.
>
> **State (1)**: Inclination anteroventral from the posterior opening to the anterior opening.
>
> **State (2)**: Less anteroventrally inclined, with an almost anteroposterior orientation close to the anterior opening.

### 3.3 Inclination of the cranial roof

Digital study of the braincase of our sample enabled observation of notable variations in the internal vault inclination (= IVI) relative to the anteroposterior axis of the cranium. The values of the IVI angle (see Material & Methods) show an interesting pattern in our sample that does not correlate (*R^2^*= 0.1768; *P-value* = 0.3246) with size (Figure 11). Whereas *Glyptodon* is distinguished from the other cingulates by an exceedingly high inclination combined with a large size, the large-sized *Vassallia* has a very low cranial roof inclination (Figure 11). *Dasypus* (adult specimen), *Stegotherium*, *Chlamyphorus*, *Proeutatus* and Miocene glyptodonts are also characterized by high inclinations (Figure 11). The juvenile *Cabassous* also displays a strong inclination, which decreases with age whereas the inclination increases with age in *Dasypus* (Figure 11). *Calyptophractus* shows an intermediate cranial roof inclination (Figure 11). The outgroups, euphractines and *Cabassous* (adult) exhibit low values, and the remaining *Peltephilus* and tolypeutines are even characterized by a negative tilting of IVI (Figure 11).

**FIGURE 11.**
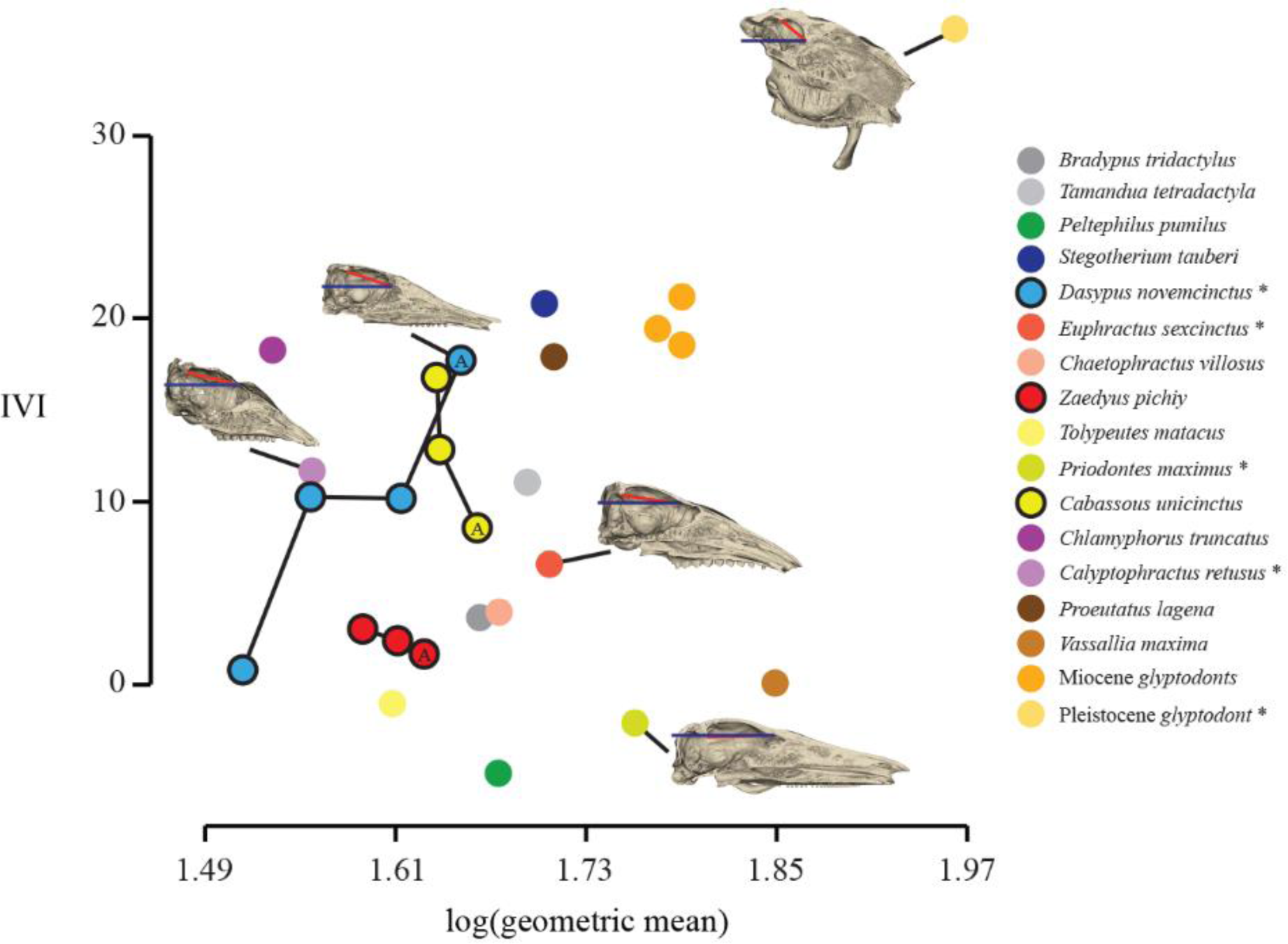
Distribution of the internal vault inclination (IVI) with respect to cranial size (= log(geometric mean)) in our sample. For the three developmental series, A symbolizes the adult specimen. Specimens marked with an asterisk are illustrated by cranial reconstructions in sagittal view in the graph. Scale = 1 cm.

We propose to treat the IVI angle as a continuous character to scrutinize evolutionary scenarios (see Table 2).

> ***Character 8*** (continuous): Internal Vault Inclination.

## 4. DISCUSSION

Due to their very unusual anatomy, even within the Cingulata, the phylogenetic placement of glyptodonts has been long debated (Burmeister, 1874; Scott, 1903; Carlini & Zurita, 2010; Fariña et al., 2013). Recent molecular analyses have proposed a new hypothesis (i.e., as sister group of the clade tolypeutines + chlamyphorines – Delsuc et al., 2016; Mitchell et al., 2016) at odds with the multiple hypotheses based on morphological analyses (Engelmann, 1985; Gaudin & Wible, 2006; Billet et al., 2011; Herrera et al., 2017). The recent investigation of the internal anatomy in the *Dasypus novemcinctus* species complex (Billet et al., 2017) underlined the potential phylogenetic signal hidden in the internal cranial structures of cingulates. Comparative studies on the paranasal sinuses and on brain endocasts of cingulates have been performed, but could not provide consistent information on the placement of glyptodonts within the Cingulata (Zurita et al., 2011; Fernicola et al., 2012; Tambusso & Fariña, 2015a). Our comparative study complements these previous efforts by adding the analysis of the dental alveolar cavities (previously only explored in the glyptodont genus *Eucinepeltus* by González Ruiz et al., 2020) and of various canals involved in the vascularization and innervation of the cranium (see Wible & Gaudin, 2004). Based on an extensive sample of extant and extinct cingulates, our survey enabled us to describe many new aspects of the internal cranial anatomy in the group, and to propose 8 potential characters with helpful phylogenetic information to aid in determining the placement of glyptodonts within Cingulata.

The anatomical variation of the dental alveolar cavities in cingulates comprises many aspects including the height, curvature, and orientation of teeth, along with their number and distribution on the rostrum (Vizcaíno, 2009). Among our observations, we have proposed two characters with potential bearing on the affinities of glyptodonts. The first corresponds to the position of the most dorsal point of these cavities in lateral view (character 1). The dorsal margins of the alveoli form a dorsally convex line for the whole dental row in most cingulates, except in taxa with reduced teeth. This most dorsal point is situated posteriorly (i.e., at Mf5-6) in *Prozaedyus*, *Zaedyus*, chlamyphorines, pampatheres and glyptodonts in our sample. The distribution of this character would be congruent with the hypothesis of a close relationship between pampatheres and glyptodonts, as suggested by many previous authors (Patterson & Pascual, 1972; Gaudin & Wible, 2006; Billet et al., 2011; Herrera et al., 2017), but also supports the clade formed by the latter two with chlamyphorines, as proposed by the morphological analysis of Mitchell et al. (2016) constrained by the molecular backbone (Figure 12). On the other hand, *Proeutatus*, generally considered to be a close relative of the clade pampatheres + glyptodonts (Engelmann, 1985; Gaudin & Wible, 2006; Billet et al., 2011; Gaudin & Lyon, 2017), has its most dorsal point situated much further anteriorly (i.e., *ca.* Mf1-3).

**FIGURE 12.**
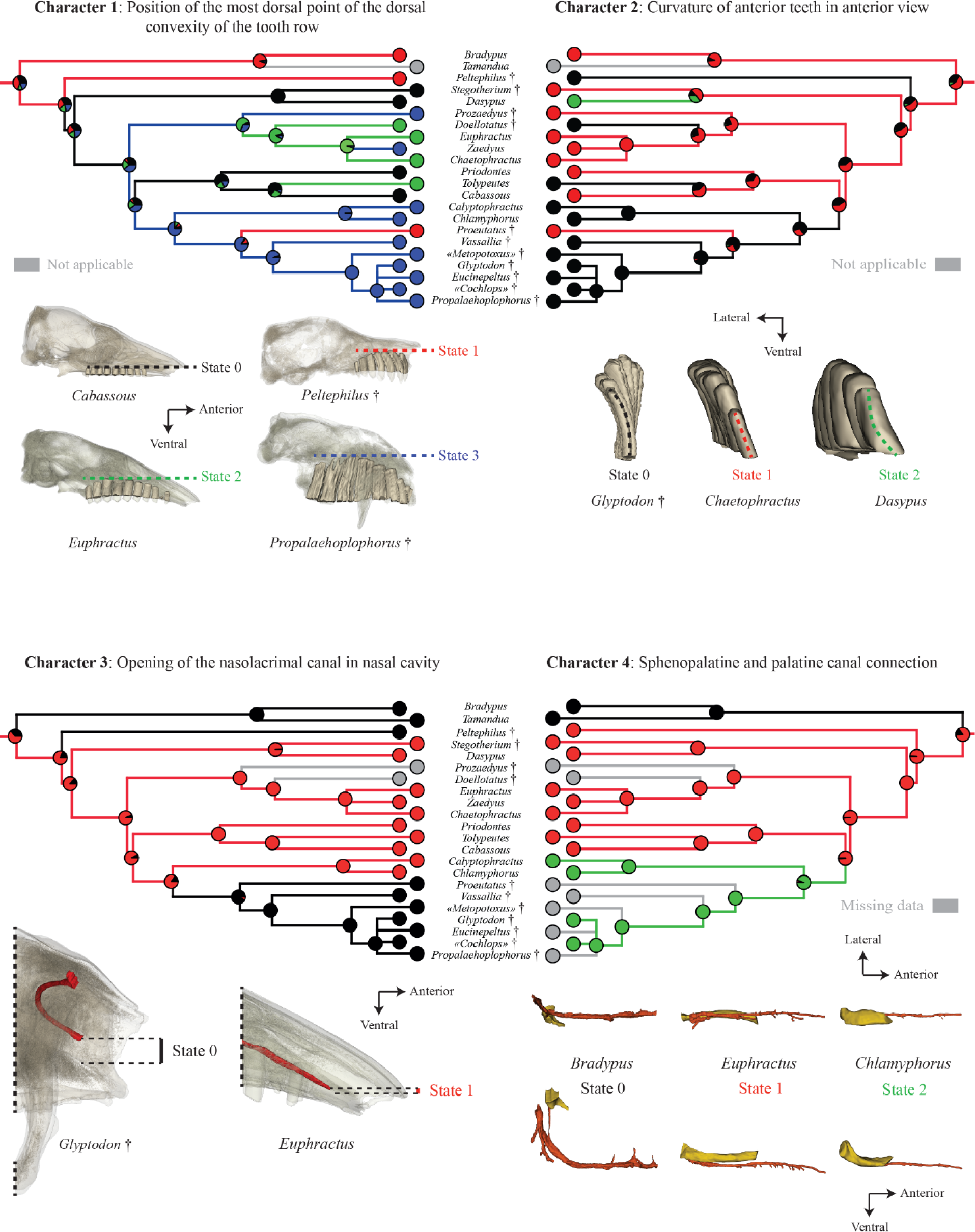
Reconstructed evolutionary scenarios for the endocranial characters 1 – 4 plotted on the reference cladogram with ancestral state estimation for internal nodes (see Materials & Methods, Supporting information 1 and Figure S3). Each character is illustrated, and the relationship between color and coding is indicated within the figure itself. † represents extinct genera. Scale = 1 cm.

The second character derived from dental alveolar cavities corresponds to the curvature of the anterior teeth in anterior view (character 2). As for character 1, chlamyphorines and pampatheres resemble glyptodonts in this aspect, exhibiting an inward curvature. *Peltephilus*, *Doellotatus* and also *Tolypeutes* show a similar condition (Figure 12). As shown by the reconstructed evolutionary scenario (Figure 12), this character could still support the morphological hypothesis of Mitchell et al. (2016) of a close relationship between chlamyphorines and glyptodonts. The distribution of this character on the baseline cladogram shows some homoplasy, however, with the condition in *Proeutatus* contrasting with that of glyptodonts, while *Peltephilus* resemble the latter (Figure 12). Another point of interest concerning the variation of dental alveolar cavities is the potential allometric pattern identified by the correlation between the relative height of the dental row and cranium size (Figure 4). This needs to be analyzed further and compared to other allometric patterns seen on the cranium. Mammals generally show an allometric elongation of the rostrum as size increases, known as craniofacial allometry (Cardini & Polly, 2013; Cardini, 2019). In *Dasypus*, this allometric pattern mostly affects the proportions of the rostrum horizontally, much less vertically (Le Verger et al., 2020). The possible allometric increase in the height of the dentition in cingulates thus seems unrelated to craniofacial allometry but this certainly requires further testing. In addition, we do not know whether allometry could affect the variation of the two aforementioned alveolar cavity characters, but the resemblance of small taxa (i.e., chlamyphorines) and large taxa (i.e., pampatheres and glyptodonts) seems to argue against this. In any case, our study brings to light interesting new variation regarding the upper tooth roots/alveolar cavities of cingulates, which have in the past been compared to one another on the basis of their number, the presence/absence of teeth in the premaxillary, tooth wear, histology, and the orientation of the long axis of the teeth relative to the long axis of the dental row (characters 1, 3 to 8 – Gaudin & Wible, 2006; see discussion for glyptodonts regarding these characters – Scott, 1903; Ferigolo, 1985; González Ruiz et al., 2015).

Our initial observations on the overall spatial organization of the alveolar cavities point towards a resemblance between chlamyphorines, pampatheres and glyptodonts, which would be partly compatible with molecular results (Delsuc et al., 2016; Mitchell et al., 2016). However, we believe further information could be gained by more detailed quantified analyses of shape to more fully reflect the complex spatial variation of the dental alveolar cavities.

Our analysis of selected intracranial canals described various aspects of their variation, which were unknown in cingulates until now. Furthermore, it enabled us to propose five characters, that provide potentially significant phylogenetic information. This analysis also demonstrated the need to further examine the variation of several structures (e.g., canal of the frontal diploic vein).

Both the trajectory and position of the nasolacrimal canal vary within the sample. A trajectory that is first directed medially at its posterior opening (i.e., the lacrimal foramen) represents a unique characteristic of glyptodonts within the Cingulata (Figures 6 & 7). The most informative feature on this canal regarding the relationships of glyptodonts with other cingulates is the relative height of its anterior opening within the nasal cavity (character 3 – Figure 12). In most cingulates, this opening is positioned ventrally in the nasal cavity, close to the hard palate, whereas in glyptodonts it is near the vertical midpoint of the cavity. The distribution of this character on the baseline cladogram supports the node linking *Proeutatus*, pampatheres and glyptodonts (Figure 12), as in several previous morphological analyses (Engelmann, 1985; Gaudin & Wible, 2006; Billet et al., 2011; Gaudin & Lyon, 2017).

The course and connection between the sphenopalatine canal and the palatine canal is characterized by a complex pattern which is very peculiar in glyptodonts (Figures S6 & S7). The degree of fusion between the two canals was variable within our sample. Because of taphonomic issues, only “*Cochlops*” and *Glyptodon* allowed us to observe a fusion of the two canals in glyptodonts. This fusion was also present as in the chlamyphorines. The uncertain condition in *Proeutatus* and pampatheres does not allow us to draw clear conclusions (character 4 – Figure 12), but the distribution of this character on the baseline cladogram could be congruent with the morphological hypothesis of Mitchell et al. (2016) (Figure 12). However, caution is warranted, because the confluence between the sphenopalatine foramen and the caudal palatine foramen is known to exhibit substantial intraspecific variation in several taxa (see character 71, Gaudin & Wible, 2006) including *Euphractus* (see Wible & Gaudin, 2004). In addition, the condition in pampatheres needs further investigation because a caudal palatine foramen has been identified in the floor of the sphenopalatine canal in *Holmesina* (see Gaudin & Lyon, 2017). Our study also reveals a strong resemblance between pampatheres and glyptodonts, which share a dorsal position of the sphenopalatine canal in relation to the nasopharyngeal canal (Figure S7). This feature could have been coded as a character, but an investigation of potential allometry for this trait is required before coding.

The course, orientation and connections of the transverse canal also vary in our sample. We were unable to identify this canal in *Tolypeutes*, chlamyphorines and *Glyptodon* as well as *Bradypus*, *Peltephilus* and *Stegotherium*. Its presence in the *Proeutatus*, pampatheres, Miocene glyptodonts and many other cingulates suggests that it has been lost in *Glyptodon*. Apart from this potential loss, the distribution on the baseline cladogram of the character corresponding to the orientation of the branch directly connected to the transverse canal foramen may support the common morphological hypothesis of a close relationship among *Proeutatus*, pampatheres and glyptodonts (character 5 – Figure 13; see also Engelmann, 1985; Gaudin & Wible, 2006; Billet et al., 2011; Gaudin & Lyon, 2017). Moreover, our work is congruent with the matrix of Gaudin & Wible (2006) regarding the presence/absence of the transverse canal foramen for most taxa (character 111) with the exceptions of *Tamandua* (coded as absent in Gaudin & Wible, 2006), *Stegotherium* (coded as present in Gaudin & Wible, 2006) and *Doellotatus*. In the case of the latter, the specimen available for our study was poorly preserved for this character (coded as present in Gaudin & Wible, 2006). Our study also allows us to confirm the hypothesis of Wible & Gaudin (2004), which suggested that the canal crosses the midline of the cranium in *Euphractus sexcinctus*. In their fetuses, they did not observe this transverse pattern, and doubted whether it existed in adult specimens (Wible & Gaudin, 2004). We can confirm this pattern in our specimen (Figure 8). This well-known circulation pattern in several marsupials is much less clear in placentals, for which the occurrence of a foramen for the transverse canal (often without knowing whether it crosses the cranium midline) is most often interpreted as a case of convergence (Sánchez-Villagra & Wible, 2002). In addition, our analysis of the youngest specimen of *Zaedyus* also suggests that the transverse canal is not formed in young euphractine individuals, in accordance with the observations of Wible & Gaudin (2004). However, in young dasypodines, we clearly observe this feature (Figure 11). For a large majority of taxa in our sample, we can confirm that a transverse canal crossing the midline is present in cingulates.

**FIGURE 13.**
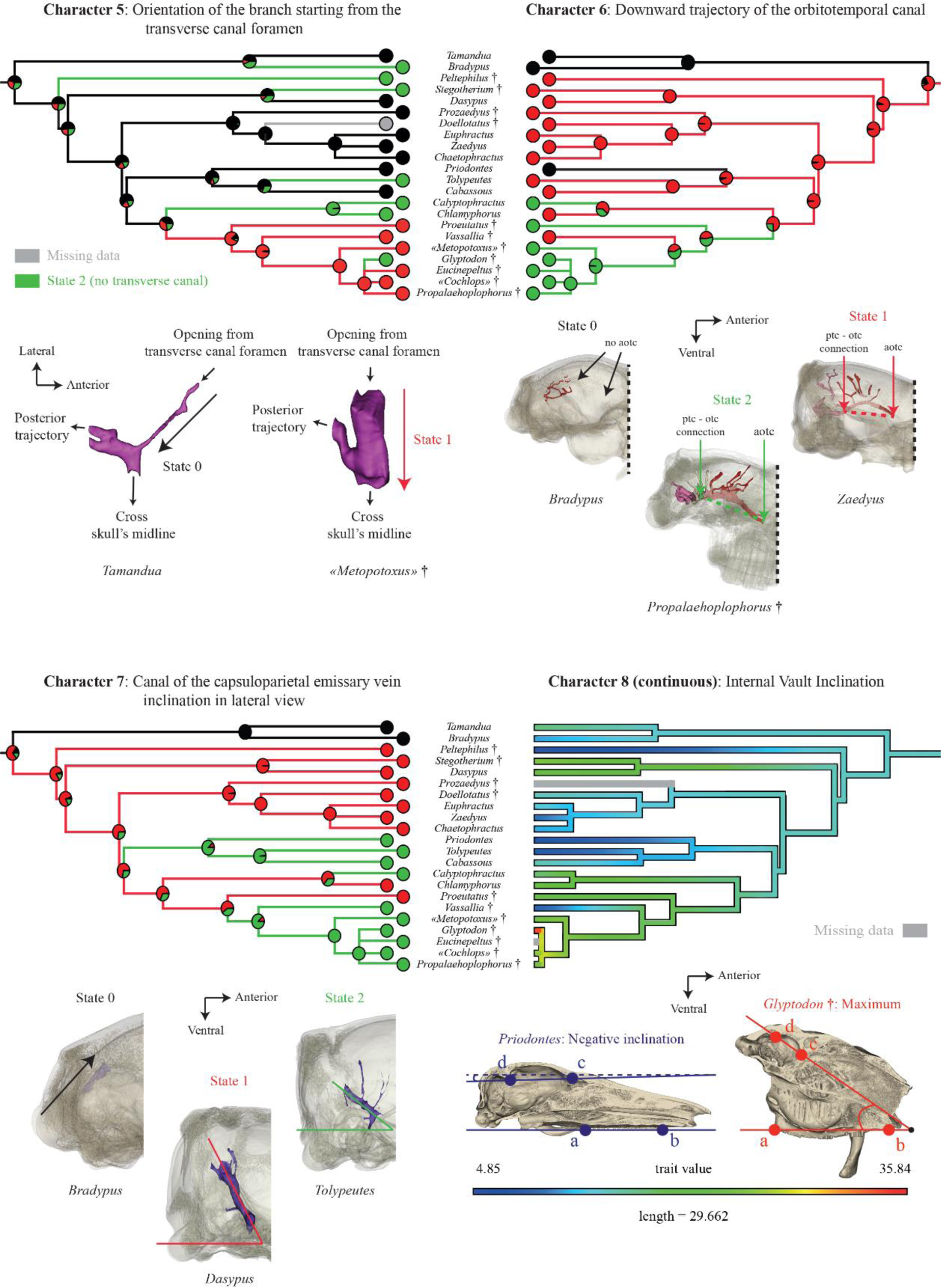
Reconstructed evolutionary scenarios for the endocranial characters 5-8 plotted on the reference cladogram with ancestral state estimation for internal nodes (see Materials & Methods, Supporting information 1 and Figure S3). Each character is illustrated and the relationship between color and coding is explained within the figure. † represents extinct genera. For character 8 (= IVI), the angle between the anteroposterior axis of the cranium (defined by the line connecting the most posterior (a) and anterior (b) edges of the tooth row) and the straight line connecting the most dorsal point of the annular ridge (c) and the most ventral point of the tentorial process (d). Scale = 1 cm.

The distribution on the baseline cladogram of the character corresponding to the anterior course of the orbitotemporal canal also provided interesting information (character 6 – Figure 13). *Calyptophractus*, *Proeutatus* and glyptodonts share a strong downward trajectory of this canal, with an anterior opening located much lower in lateral view than its posterior connection of the orbitotemporal canal to the posttemporal canal. The evolutionary scenario for this character is, however, unclear for glyptodonts and their close allies, and homoplasy seems to be present (Figure 13). Our study also highlights the presence of a foramen for the anterior opening of the orbitotemporal canal in all cingulates except *Priodontes* whereas this foramen was scored as absent in *Vassallia* and *Chlamyphorus* in Gaudin & Wible (2006, character 78), and is often unknown for other cingulate taxa. Based on our observation, there is no doubt that the canal opens in the orbitotemporal region in *Vassallia*, while this opening is described as absent in *Holmesina floridanus* Robertson, 1976 by Gaudin & Lyon (2017). The opposite discrepancy exists regarding the foramen for the frontal diploic vein, absent in *Vassallia* and described as present in *Holmesina floridanus* by Gaudin & Lyon (2017). As these two foramina are usually located relatively close to each other in the orbitotemporal region (bearing many foramina), we suggest that misidentifications may have occurred between the opening of these two canals, if they are present, or with ethmoidal foramina (Figure S4). The only way to confirm this hypothesis would be to scan Gaudin & Lyon (2017) specimens of *Holmesina floridanus*.

Another character we investigated concerned the inclination of the canal for the capsuloparietal emissary vein in lateral view (character 7 – Figure 13). All tolypeutines, *Calyptophractus*, *Vassallia* and glyptodonts share a less steep inclination than that observed in other cingulates, with a part close to the anterior opening oriented almost directly to the anterior direction. The distribution of this character could provide support for the molecular hypothesis of a closer relationship of the glyptodonts with chlamyphorines and tolypeutines (Delsuc *et al*., 2016; Mitchell *et al*., 2016; Figure 13). However, the condition is reversed in *Chlamyphorus* and *Proeutatus*, creating homoplasy that casts doubt on the evolution of this trait. The trajectory of this canal has never been scored in prior phylogenetic analyses of the Cingulata. In the literature, only two characters have been scored that indirectly concern this canal, characters 119 and 120 of Gaudin & Wible (2006), related to the presence/absence of the postglenoid foramen and suprameatal foramen, respectively. These characters deliver contrasting grouping among cingulates (Gaudin & Wible, 2006).

Finally, the internal inclination of the cranial vault relative to the anteroposterior axis of the cranium also helps identify some cingulate taxa that may resemble glyptodonts in this regard (character 8; Figure 13). The distribution of this quantitative trait on the baseline cladogram indicates that the dasypodines, chlamyphorines, *Proeutatus* and glyptodonts exhibit a strong IVI angle, whereas *Vassallia* possesses a very weak angle. This strong difference between glyptodonts and pampatheres was already highlighted by Tambusso & Fariña (2015b). The variation of this trait was not explained by allometry in our sample. A highly inclined vault may have been evolved independently in the dasypodines and in glyptodonts and their allies. Its distribution could provide support for a relationship among chlamyphorines, *Proeutatus* and glyptodonts to the exclusion of pampatheres or the latter may have undergone a reversal (Figure 13). The vault inclination in these taxa seems to be partly congruent with the distribution of a character proposed by Billet et al. (2011) and discussed by Delsuc et al. (2016), i.e., the dorsal position of the ventral surface of the auditory region relative to the palate (character 78 – Billet et al., 2011). This is a characteristic of chlamyphorines, eutatines, pampatheres, and glyptodonts in Billet et al.’s (2011) analysis. The potential relationship of these two aspects of the braincase should be tested in future studies.

In conclusion, our investigation of the cranial canals and alveolar cavities highlighted several endocranial characters that support a resemblance of glyptodonts with chlamyphorines, *Proeutatus* and pampatheres. Except for character 7, our investigation did not bring to light new, and strong resemblances between tolypeutines and glyptodonts that would support their close relationship as suggested by molecular studies (Delsuc et al., 2016; Mitchell et al., 2016). In short, our study of internal anatomy lends further credence to recent reports of a greater morphological resemblance between glyptodonts and chlamyphorines than tolypeutines (Delsuc et al., 2016; Mitchell et al., 2016). In this regard, it is worth noting that the chlamyphorines + tolypeutines relationship received distinctly lower statistical support than most other clades in mitogenomic analyses of cingulate phylogeny (Delsuc et al., 2016; Mitchell et al., 2016).

The congruence of these new internal characters with past morphological phylogenetic analyses of the group is difficult to evaluate since the results of the latter have been rather unstable (e.g., Engelmann, 1985; Abrantes & Bergqvist, 2006; Gaudin & Wible, 2006; Billet et al., 2011). However, a close resemblance of glyptodonts to the eutatine *Proeutatus* and the pampatheres is supported by several characters in our study, which would be congruent with several recent phylogenetic hypotheses based on morphological data (Engelmann, 1985; Gaudin & Wible, 2006; Billet et al., 2011; Gaudin & Lyon, 2017). More generally, internal cranial anatomy seems to show similarities among chlamyphorines, *Proeutatus*, pampatheres, and glyptodonts (see also Tambusso et al., 2021), but new analyses are needed that incorporate this new data before firmer phylogenetic conclusions can be reached. Our study provides a non-exhaustive account of internal canals of the cranium. Internal structures still need to be studied to select new characters in particular for endocranial traits that we briefly describe but do not explore in detail. This is notably the case of the canal for the frontal diploic vein or the relative position of the sphenopalatine canal in relation with the nasopharyngeal canal, whose patterns of variation remain unclear at the moment. Another promising way forward was also revealed by our developmental series in which *Cabassous* varies in a manner different from that of *Dasypus* and *Zaedyus* as far as the inclination of the cranial roof is concerned (Figure 11). A better exploration of the developmental series with more specimens per species and more species sampled could provide a better understanding of their ontogenetic trajectories, which can be coded for phylogenetic analyses (e.g., Bardin et al., 2017). Further studies are also needed to better understand the patterns of variation of the internal characters highlighted here, which in some respects may be affected by allometry as we have highlighted for alveolar cavities and the canal for the capsuloparietal emissary vein.

## ACKNOWLEDGMENTS

We are grateful to Christiane Denys, Violaine Nicolas, and Géraldine Véron, (Muséum National d’Histoire Naturelle, Paris, France), Roberto Portela Miguez, Louise Tomsett, and Laura Balcells (British Museum of Natural History, London, UK), Neil Duncan, Eileen Westwig, Eleanor Hoeger, Ross MacPhee, Marisa Surovy and Morgan Hill Chase (American Museum of Natural History, New York, USA), Nicole Edmison and Chris Helgen (National Museum of Natural History, Washington, DC, USA), Jake Esselstyn (Louisiana State University, Museum of Natural Sciences, Baton Rouge, USA), Manuel Ruedi (Muséum d’Histoire Naturelle, Geneva, Switzerland), Pepijn Kamminga, Arjen Speksnijder and Rob Langelaan (Naturalis Biodiversity Center, Leiden, Holland), April Isch and Zhe-Xi Luo (University of Chicago, Chicago, USA), Adrienne Stroup and Bill Simpson (Field Museum of Natural History, Chicago, USA), Steffen Bock, Christiane Funk, Frieder Mayer, Anna Rosemann Lisa Jansen, Kristin Mahlow, Johannes Müller and Eli Amson (Museum für Naturkunde, Berlin, Germany), and Camille Grohé (University of Poitiers, Poitiers, France) for access to comparative material and/or to CT-scans. We would like to warmly thank Daniel Brinkmann (Yale Peabody Museum, New Haven, USA) for his precious help in organizing KLV’s visit to the collections of the Yale Peabody Museum. Many thanks also to Sergio Ferreira-Cardoso (Institut des Sciences de l’Evolution) for his help during the trips of KLV for data acquisition. We thank Benoit de Thoisy (Institut Pasteur de la Guyane) and Clara Belfiore for their help with the data acquisition, Olivia Plateau (University of Fribourg, Switzerland) for her help on the R script and Cyril Le Verger for his help with segmentation. We thank Renaud Lebrun (Institut des Sciences de l’Evolution), Farah Ahmed (British Museum of Natural History), Miguel García-Sanz, Marta Bellato, Nathalie Poulet and Florent Goussard (Platform AST-RX – Muséum National d’Histoire Naturelle) who generously provided help with CT-scanning. Some of the experiments were performed using the µ-CT facilities of the Montpellier Rio Imaging (MRI) platform of the LabEx CeMEB. KLV acknowledges the financial support provided by the Sorbonne Universities/ED227 transhumance international grant program. Finally, we would like to warmly thank Christian de Muizon (Centre de Recherche en Paléontologie – Paris), Timothy Gaudin (University of Tennessee), Robert J. Asher (University of Cambridge), Sophie Montuire (University of Bourgogne), Isabelle Rouget (Centre de Recherche en Paléontologie – Paris), Allowen Evin and Lionel Hautier (Institut des Sciences de l’Evolution) for reviewing a previous version of this work in the PhD dissertation of KLV.

## AUTHOR CONTRIBUTIONS

KLV contributed to data acquisition, data analyses/interpretations, drafting of the manuscript, and study design. LRG contributed to data acquisition and critical revision of the manuscript. GB contributed to data acquisition, study design, and critical revision of the manuscript.

## SUPPLEMENTARY DATA

SUPPORTING INFORMATION 1: Matrix and Analytical Parameters.

For de description of each characters and the previous coded taxa see Billet *et al*. (2011).

### Parameters

- Dimensions ntax = 22 nchar = 125;

- Format datatype = standard;

- Gap = -;

- Missing = ?;

- States = “0 1 2 3 4 5 6”.

- Heuristic search settings: Optimality criterion = parsimony;

- Character-status summary: Of 125 total characters: 27 characters are of type ‘ord’ (Wagner); 98 characters are of type ‘unord’; All characters have equal weight; All characters are parsimony-informative; Gaps are treated as “missing”; Multistate taxa interpreted as polymorphism.

- Starting tree(s) obtained via stepwise addition;

- Addition sequence: random;

- Number of replicates = 1000;

- Starting seed = generated automatically;

- Number of trees held at each step = 10;

- Branch-swapping algorithm: tree-bisection-reconnection (TBR) with reconnection limit = 8;

- Steepest descent option not in effect;

- Initial ‘Maxtrees’ setting = 100;

- Branches collapsed (creating polytomies) if maximum branch length is zero;

- ‘MulTrees’ option in effect;

- Keeping only trees compatible with constraint-tree “kevin”;

- Trees are unrooted.

New coded taxa

#### Calyptophractus retusus

3210? 1???? 10611 12020 10012 01211 11010 12112 ??011 10000 0000? 00010 10221 11121 01011 111?0 00110 10111 110?0 0?10? 01011 2300? ???01 10101 10110

#### “Cochlops” debilis

3?1?? 2????????? ???0? ??201 ?0010 13113 ?0021 10000 0?00? 110?? ??121 21121 01111 111?1 01100 10210 11010 120-? 001?? 0??1? ????1 1211? (01)0111

#### Eucinepeltus complicatus

3211? 21101 206???0?0? 10201 10010 13113 ?0021 1?000 0?0?? 110?? 1?122 21121 00111 111?1 011?0 10210 1???0001?? ???1? ????1 12111 10111

#### Glyptodon sp.

3211? 21101 20601 02321 10?0? 10201 10010 13113 10021 10000 0?0?? 110?? 1?121 21121 00111 111?1 0?1?0 10210 11010 120?1 001?? 0??1? ???01 12111 10111

#### “Metopotoxus” anceps

3???? 2????????? ???0? ??20? ?0010 13??? ?0011 10000 ??00? 110?? ???21 21121 01111 111?1 1?1?? ?0210????? 0????????1 1211? 0011?

---

**TABLE S1.** List of adult specimens used for assessing the intraspecific variation in *Dasypus*, *Zaedyus* and *Cabassous*.

| Family/Subfamily | Species | Institutional number | Locality |
| --- | --- | --- | --- |
| Dasypodinae | *Dasybus novemcinctus* | AMNH 255866 | Bolivia, Beni, Cercado |
| Dasypodinae | *Dasybus novemcinctus* | AMNH 211668 | Bolivia, Beni, Cercado, ca. 23 kilometers west of San Javier |
| Dasypodinae | *Dasybus novemcinctus* | AMNH 262658 | Bolivia, Pando, Nicolas Suarez, 5 kilometers upstream from Cachuela |
| Dasypodinae | *Dasybus novemcinctus* | AMNH 263287 | Bolivia, Santa Cruz, Andres Ibanez, Ayacucho |
| Dasypodinae | *Dasybus novemcinctus* | AMNH 205727 | Uruguay, Lavalleja, Zapican, 12 kilometers west southwest of Zapican |
| Euphractinae | *Zaedyus pichiy* | FMNH 23810 | Argentina, Chubut |
| Euphractinae | *Zaedyus pichiy* | AMNH 94327 | Argentina, Chubut, Sarmiento, Colhue Huapi Lake |
| Euphractinae | *Zaedyus pichiy* | MNHN 1883-158 | Argentina, Santa Cruz |
| Euphractinae | *Zaedyus pichiy* | ZMB 46103 | South America, Argentina, Puerto Jenkins, -47,75779, -65,90836 |
| Euphractinae | *Zaedyus pichiy* | ZMB 46104 | South America, Argentina, Oso Marino, -47,9237, -65,83145 |
| Tolypeutinae | *Cabassous unicinctus* | MNHN 1999-1044 | French Guiana |
| Tolypeutinae | *Cabassous unicinctus* | MNHN 1998-2255 | French Guiana, Petit-Saut |
| Tolypeutinae | *Cabassous unicinctus* | AMNH 137196 | South America, Brazil, Para, Tapajos River, Igarape Brabo |
| Tolypeutinae | *Cabassous unicinctus* | AMNH 74113 | South America, Peru, Loreto, Maynas |
| Tolypeutinae | *Cabassous unicinctus* | MVZ 155192 | Peru, Rio Cenepa |

**TABLE S2.**
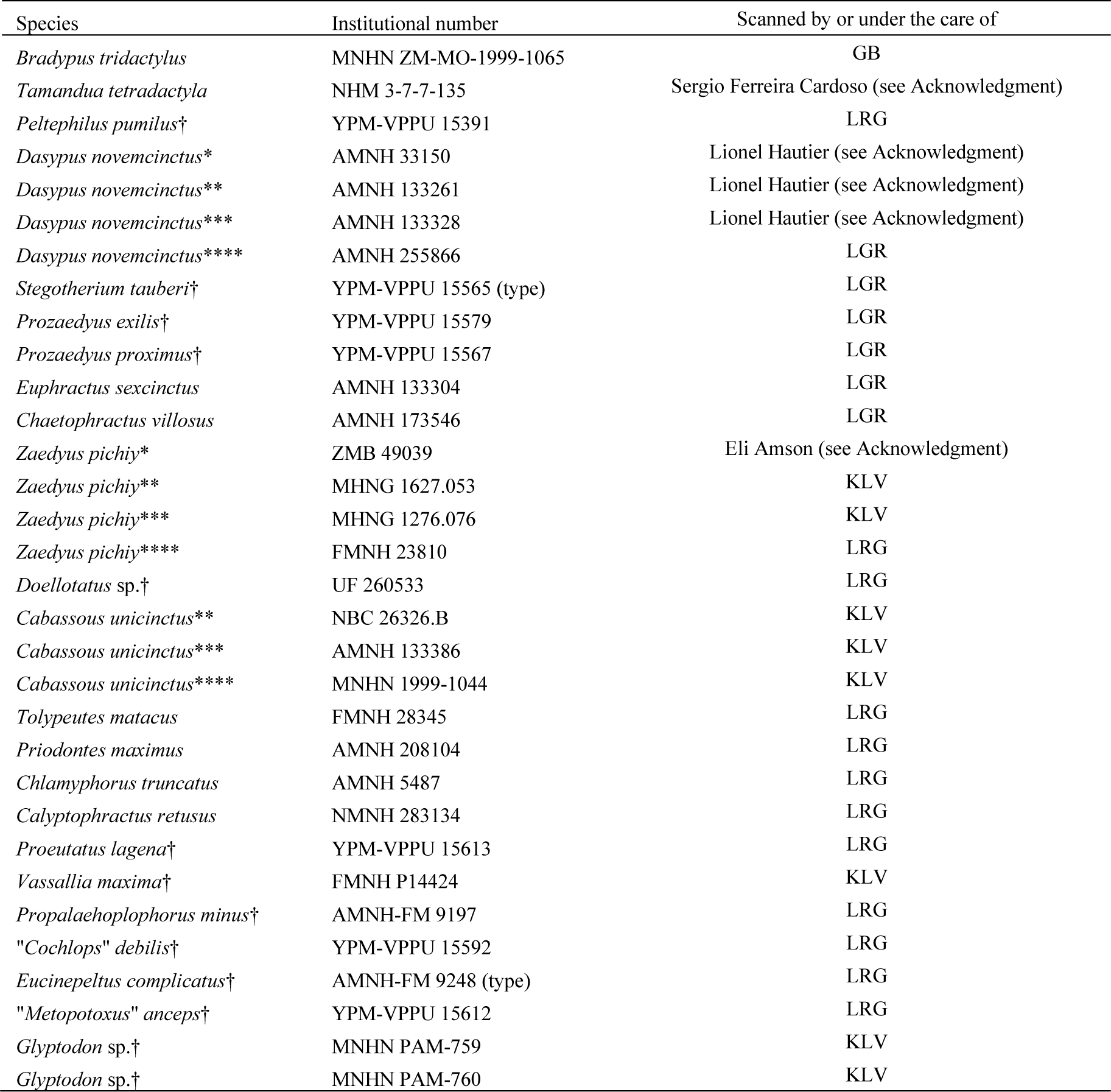
List of scanned specimens available online or under request (see Materials & Methods). Symbol: †, extinct species; *, perinatal stage; **, juvenile; ***, subadult; ****, adult; (i), author of the genus.

**TABLE S3.** List of specimens with specific dental formula. Symbol: †, extinct species; *, perinatal stage; **, juvenile; ***, subadult; ****, adult; (i), author of the genus; C, canine; dPM, decidious premolar; M, molar; Mf, molariform; Mfp, molariform in premaxillary; PM, premolar; pmx, premaxilla; Ust, unfunctional supernumary teeth.

| Family/Subfamily | Species | Institutional number | Dental formula |
| --- | --- | --- | --- |
| Bradypodidae | *Bradypus tridactylus* | MNHN ZM-MO-1999-1065 | 1 C + 4 Mf |
| Myrmecophagidae | *Tamandua tetradactyla* | NHM 3-7-7-135 | no teeth |
| Peltephilidae | *Peltephilus pumilus*† | YPM-VPPU 15391 | 1 Mfp + 6 Mf |
| Dasypodinae | *Dasypus novemcinctus* * | AMNH 33150 | 7 dPM |
| Dasypodinae | *Dasypus novemcinctus* ** | AMNH 133261 | 7 dPM |
| Dasypodinae | *Dasypus novemcinctus* *** | AMNH 133328 | 7 dPM/PM + 1 M |
| Dasypodinae | *Dasypus novemcinctus* **** | AMNH 255866 | 7 PM + 1 M |
| Dasypodidae | *Stegotherium tauberi*† | YPM-VPPU 15565 (type) | 6 Mf |
| Euphractinae | *Prozaedus exilis*† | YPM-VPPU 15579 | 8 Mf |
| Euphractinae | *Prozaedus proximus*† | YPM-VPPU 15567 | 8 Mf |
| Euphractinae | *Euphractus sexcinctus* | AMNH 133304 | 1 Mfp + 1 Ust + 8 Mf |
| Euphractinae | *Chaetophractus villosus* | AMNH 173546 | 1 Mfp + 1 Ust + 8 Mf |
| Euphractinae | *Zaedyus pichiy* * | ZMB 49039 | 6 Mf |
| Euphractinae | *Zaedyus pichiy* ** | MHNG 1627.053 | 8 Mf |
| Euphractinae | *Zaedyus pichiy* *** | MHNG 1276.076 | 8 Mf |
| Euphractinae | *Zaedyus pichiy* **** | FMNH 23810 | 8 Mf |
| Euphractinae | *Doellotatus* sp.† | UF 260533 | Incomplete specimen bearing 7 Mf (9 Mf in Gaudin & Wible, 2006) |
| Tolypeutinae | *Cabassous unicinctus* ** | NBC 26326.B | 9 Mf |
| Tolypeutinae | *Cabassous unicinctus* *** | AMNH 133386 | 9 Mf |
| Tolypeutinae | *Cabassous unicinctus* **** | MNHN 1999-1044 | 9 Mf |
| Tolypeutinae | *Tolypeutes matacus* | FMNH 28345 | 1 Ust on pmx + 9 Mf |
| Tolypeutinae | *Priodontes maximus* | AMNH 208104 | 19 Mf |
| Chlamyphorinae | *Chlamyphorus truncatus* | AMNH 5487 | 8 Mf |
| Chlamyphorinae | *Calyptophractus retusus* | NMNH 283134 | 8 Mf |
| Eutatini? | *Proeutatus lagna*† | YPM-VPPU 15613 | 1 Mfp + 8 Mf |
| Pampatheriinae | *Vassallia maxima*† | FMNH P14424 | 1 Mfp + 8 Mf |
| Glyptodontinae | *Propalaeohoplophorus minus*† | AMNH-FM 9197 | 8 Mf |
| Glyptodontinae | *"Cochlops" debilis*† | YPM-VPPU 15592 | Incomplete specimen bearing 7 Mf (8 Mf in Scott, 1903) |
| Glyptodontinae | *Eucinepeltus complicatus*† | AMNH-FM 9248 (type) | 8 Mf |
| Glyptodontinae | *"Metopotoxus" anceps*† | YPM-VPPU 15612 | Incomplete specimen bearing 7 Mf (8 Mf in Scott, 1903) |
| Glyptodontinae | *Glyptodon* sp.† | MNHN PAM-759 | 8 Mf |
| Glyptodontinae | *Glyptodon* sp.† | MNHN PAM-760 | 8 Mf |

**TABLE S4.**
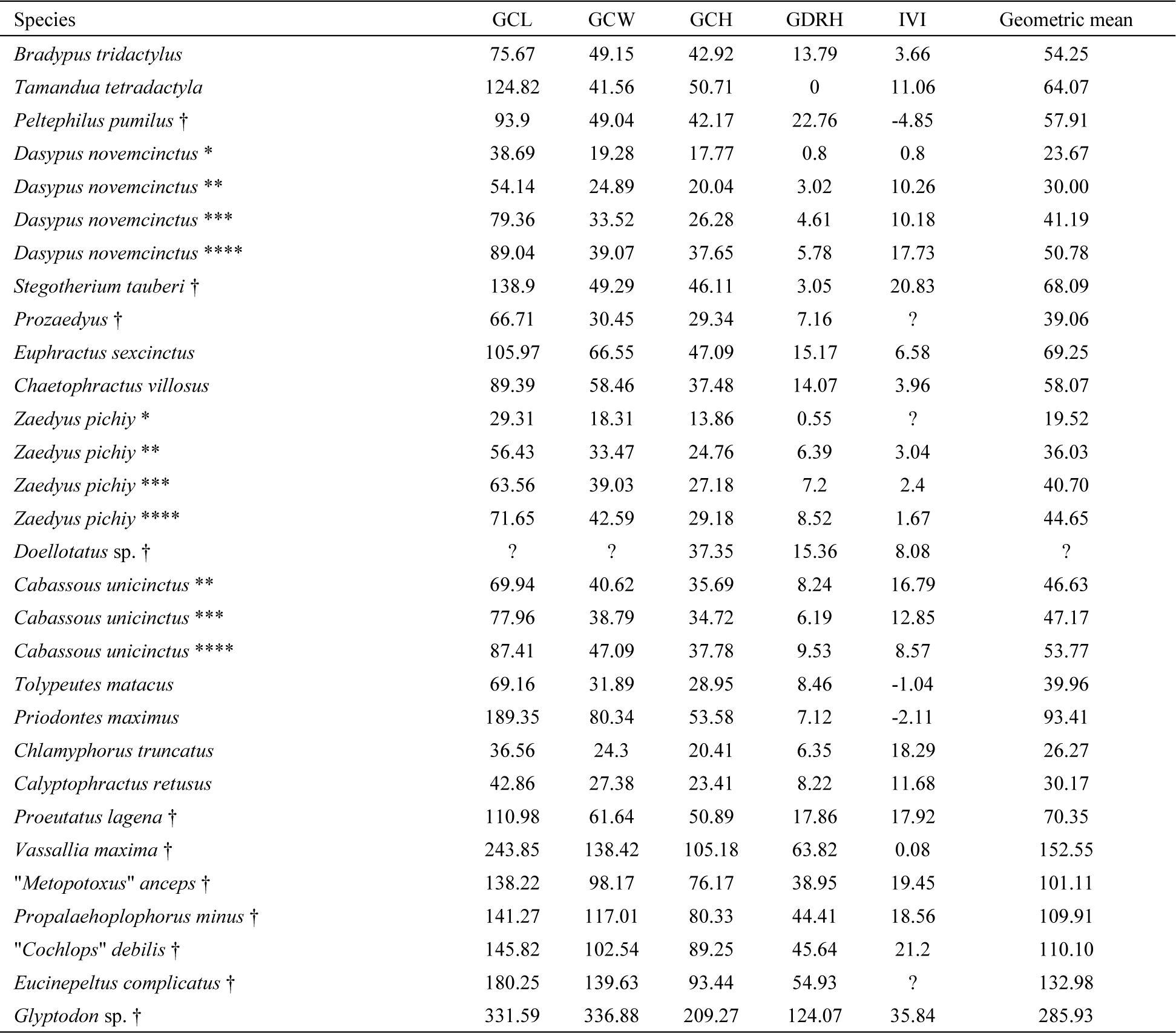
Cranial measurements in mm for GCL, GCW, GCH, GDRH and Geometric mean, and in degree for IVI (see Figure S2). Symbol: ?, missing data; †, extinct species; *, perinatal stage; **, juvenile; ***, subadult; ****, adult.

**FIGURE S1.**
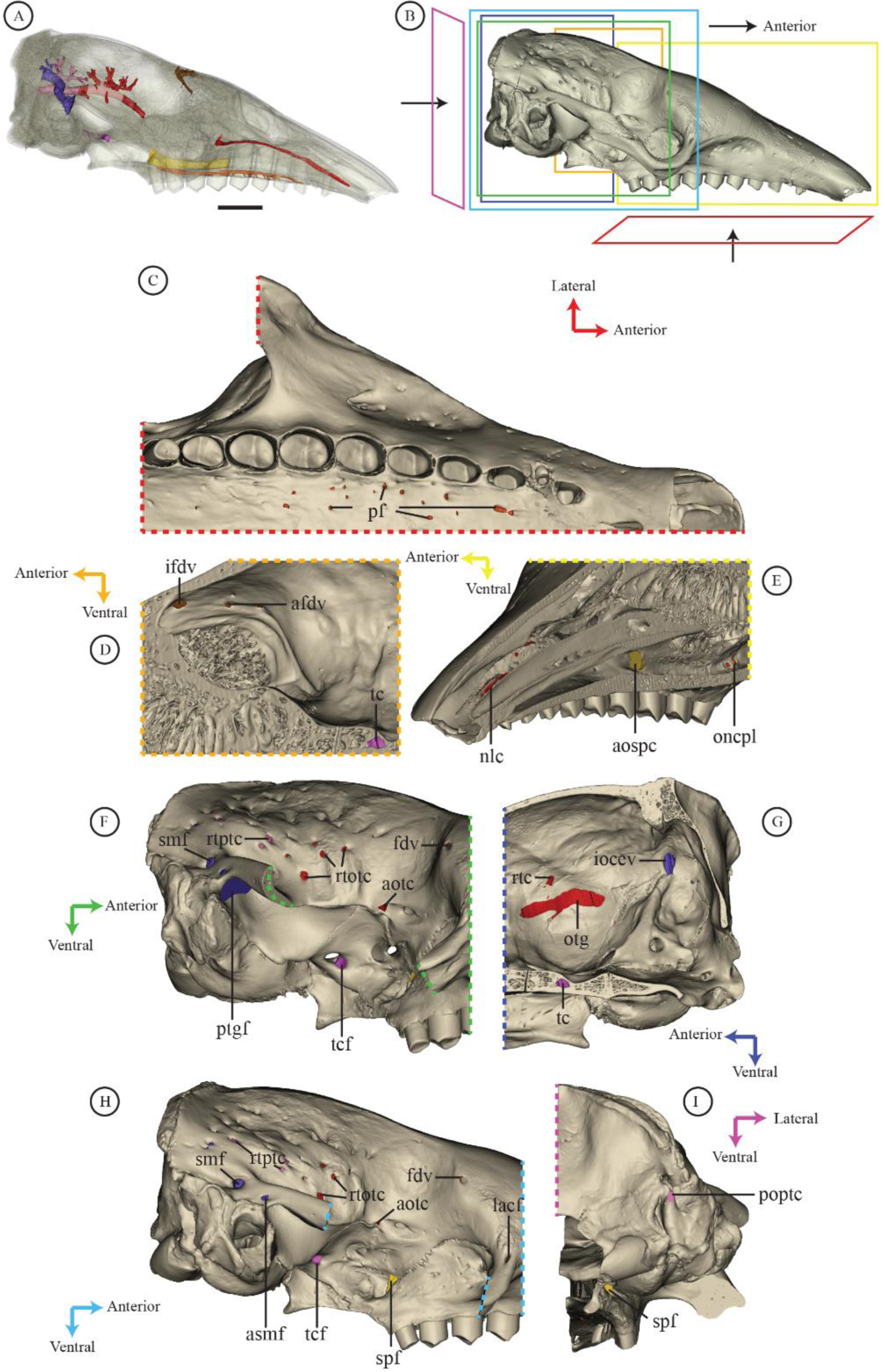
Illustration of the internal and external openings of each selected canal on a specimen of *Euphractus sexcinctus* (AMNH 133304) in transparent and non-transparent views. A, B, F and H, lateral view; C, ventral view; D and E, oblique internal view; G, lateral view of the braincase inner part; I, occipital view. Colors of the canals follow Figure 1. Abbreviations: afdv, accessory frontal diploic vein internal opening; aospc, anterior opening of the sphenopalatine canal; aotc, anterior opening of the orbitotemporal canal, asmf, accessory suprameatal foramen; fdv, foramen for the frontal diploic vein; ifdv, internal foramen for the frontal diploic vein; iocev, internal opening of the capsuloparietal emissary vein canal; lacf, lacrimal foramen; nlc, nasolacrimal canal; oncpl, opening in nasopharyngeal canal of palatine canal; otg, orbitotemporal groove; pf, palatine foramina; poptc, posterior opening of the posttemporal canal; ptgf, postglenoid foramen; ramus temporalis canal; rtotc, rami temporales foramina from the orbitotemporal canal; rtptc, rami temporales foramina from the posttemporal canal; smf, suprameatal foramen; spf, sphenopalatine foramen; tc, tranverse canal; tcf, tranverse canal foramen. Scale = 1 cm.

**FIGURE S2.**
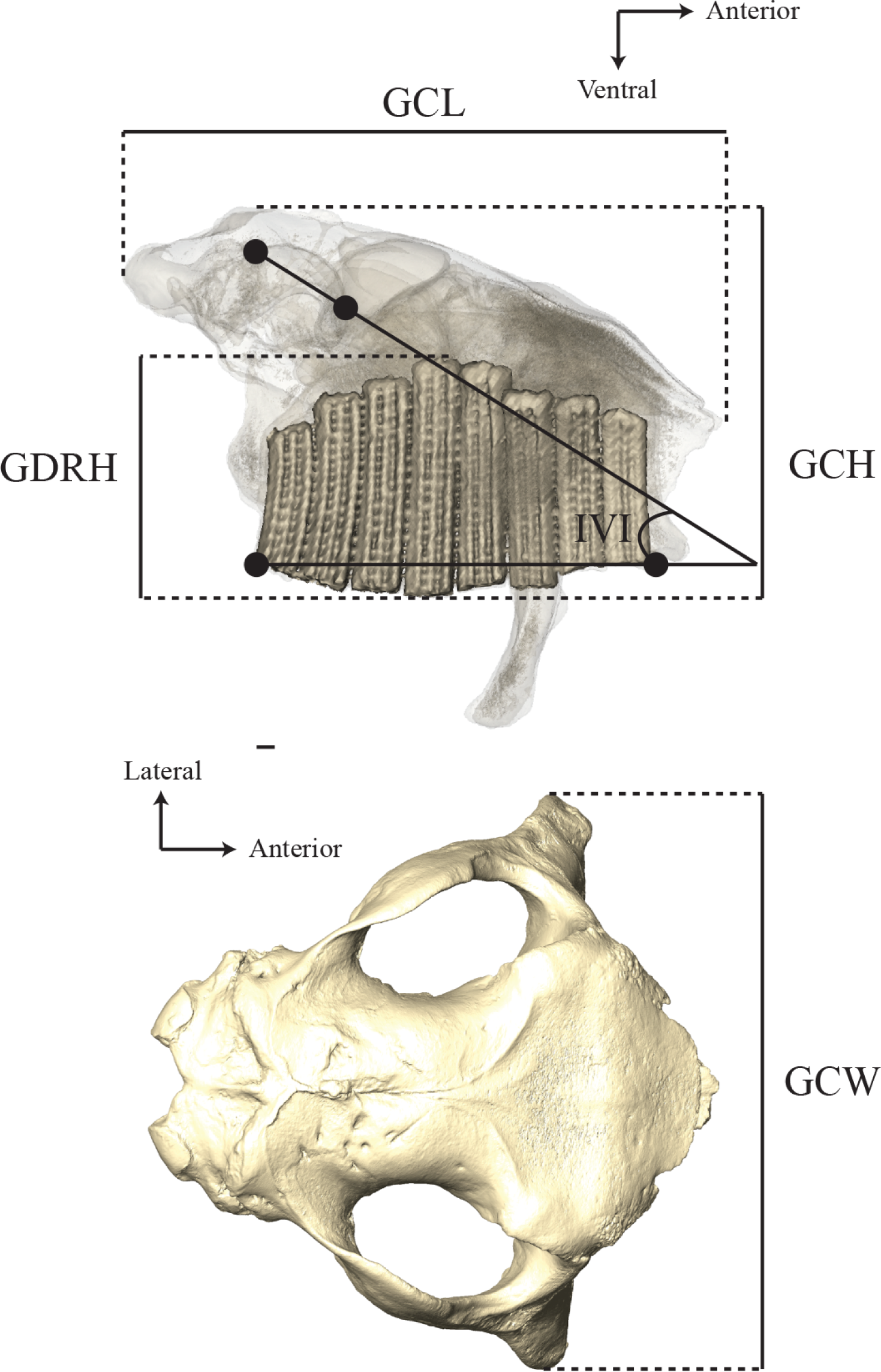
Cranial measurements (Table S2) illustrated on a transparent cranium in lateral (top) and dorsal (bottom) view of Glyptodon sp. MNHN.F.PAM 760. Abbreviations: GDRH, greatest dental row height; GCH, greatest cranium height; GCL, greatest cranium length; GCW, greatest cranium width; IVI, internal vault inclination (= angle between the anteroposterior axis of the cranium (defined by the line connecting the most anterior and posterior edges of the dental row) and the straight line connecting the most dorsal point of the annular ring and the most ventral point of the tentorial process). Scale = 1 cm.

**FIGURE S3.**
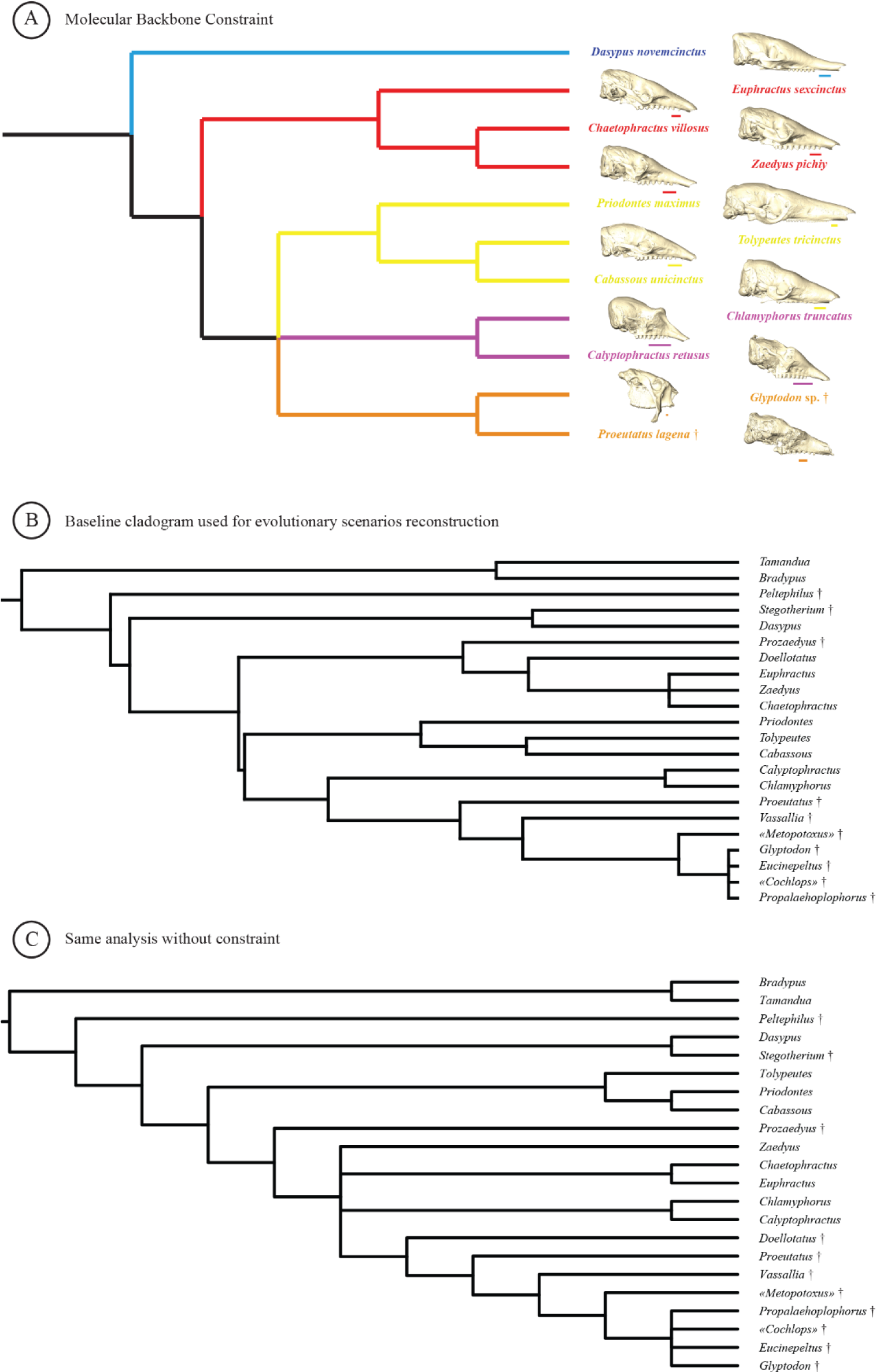
A, modified backbone constraint from Mitchell et al. (2016) used for our phylogenetic analysis (see text). *Proeutatus* is enforced as the sister group of glytptodonts in accordance with most phylogenetic analyses using morphology (Engelmann, 1985; Gaudin & Wible, 2006; Billet et al., 2011; Herrera et al., 2017). B. Topology of the baseline cladogram. C. Topology of the strict consensus from the same analysis without constraint. See Supporting information 1 for phylogenetic analysis.

**FIGURE S4.**
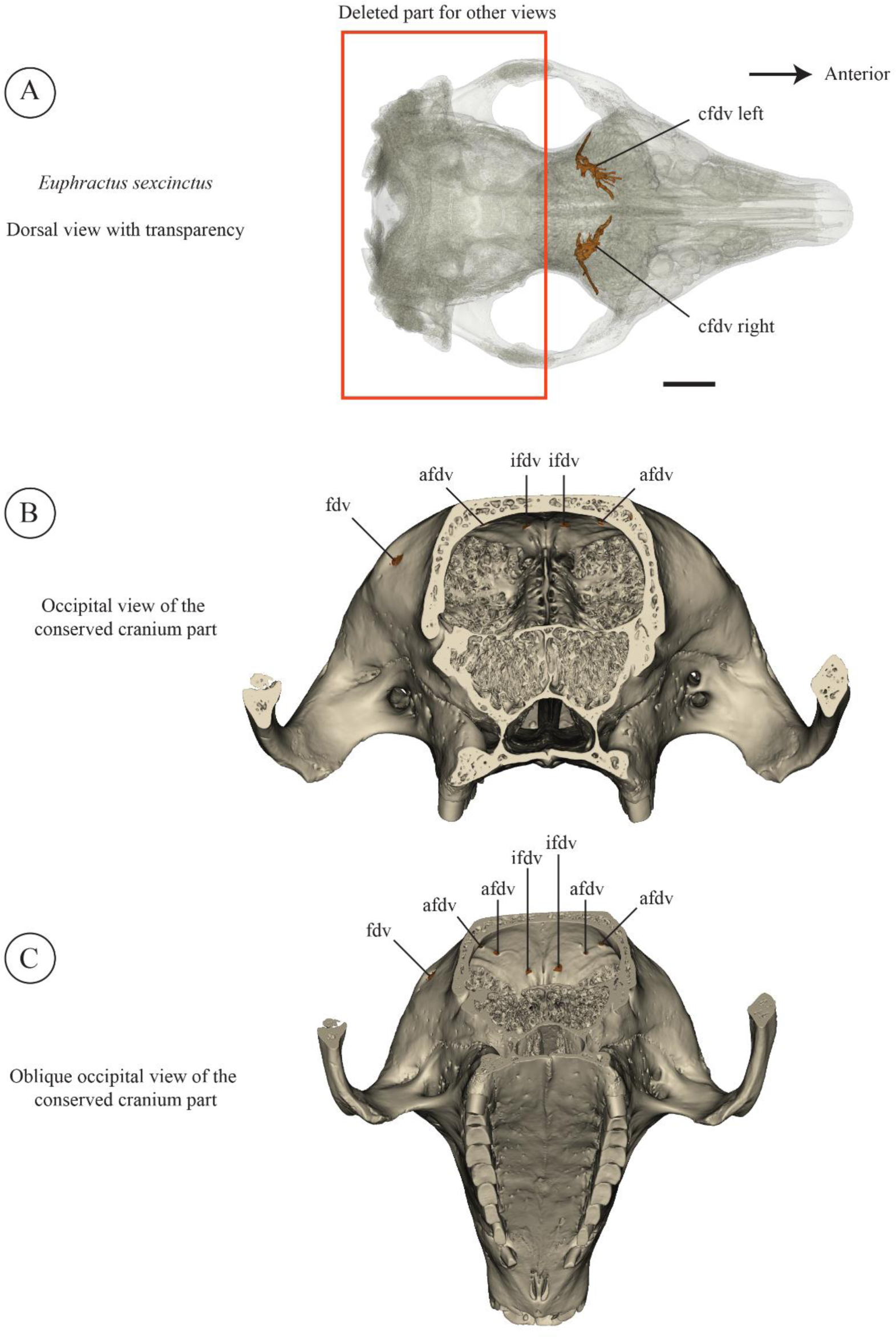
Internal/external opening and canal trajectory for the right and left frontal diploic vein in *Euphractus sexcinctus* (AMNH 133304). A. Dorsal view in transparency with the two canals for the frontal diploic vein. The two canals do not cross the cranium midline. B. Occipital view of the conserved cranium part with internal opening. C. Oblique (more ventral) occipital view to facilitate the visibility of internal openings. Colors of the canals follow Figure 1. Abbreviations: afdv, accessory frontal diploic vein internal opening; cfdv, canal for the frontal diploic vein; fdv, foramen for the frontal diploic vein; ifdv, internal foramen for the frontal diploic vein. Scale = 1 cm.

**FIGURE S5.**
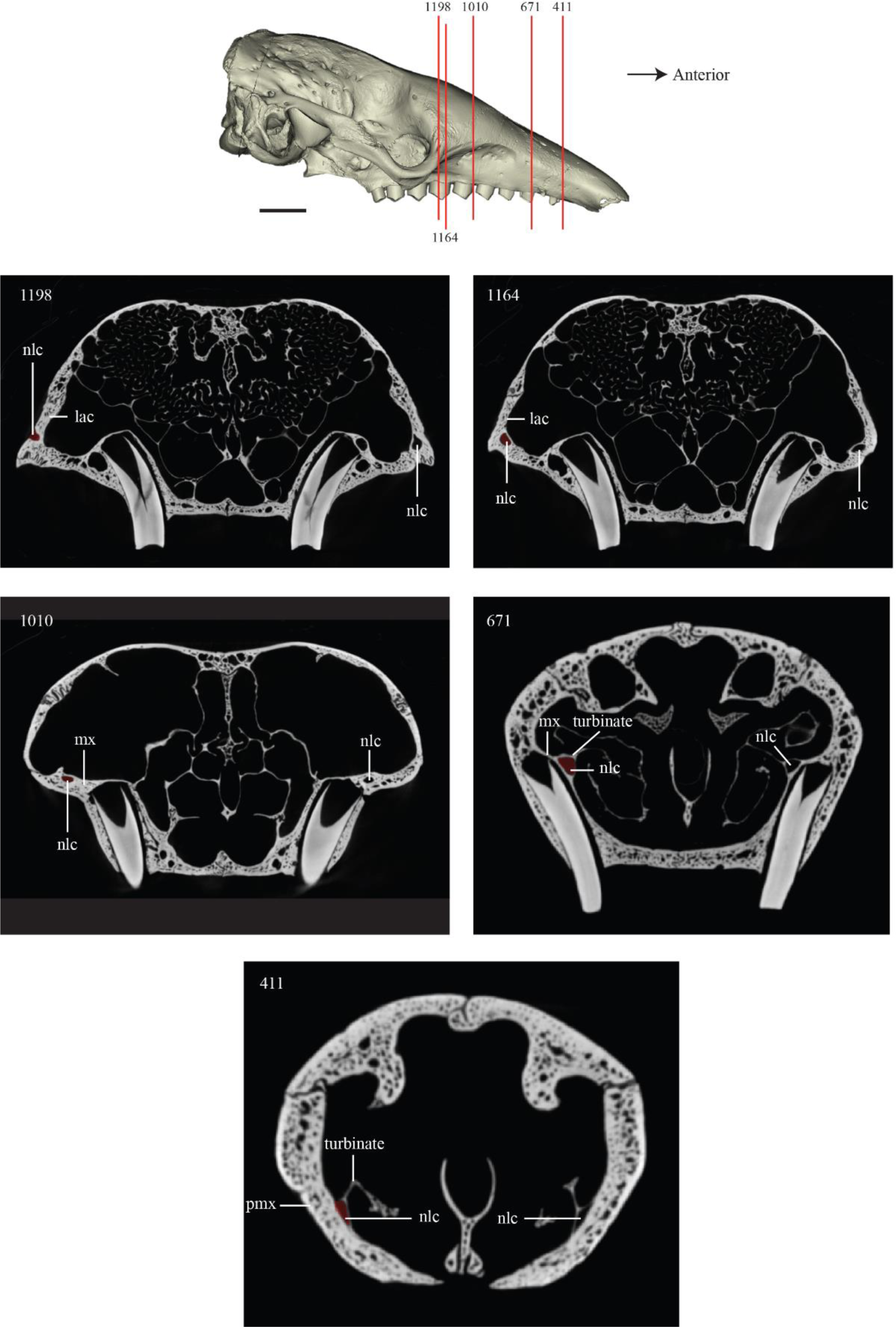
Identification of the bones enclosing the nasolacrimal canal in *Euphractus sexcinctus* (AMNH 133304) illustrated by a cranium in right lateral view (top). The slices analyzed are indicated on the cranium by their number in the image stack and illustrated below. The nasolacrimal canal is reconstructed in red on each slice, on the left side of the image. Abbreviations: lac, lacrimal; mx, maxillary; nlc, nasolacrimal canal; pmx, premaxillary. Scale = 1 cm.

**FIGURE S6.**
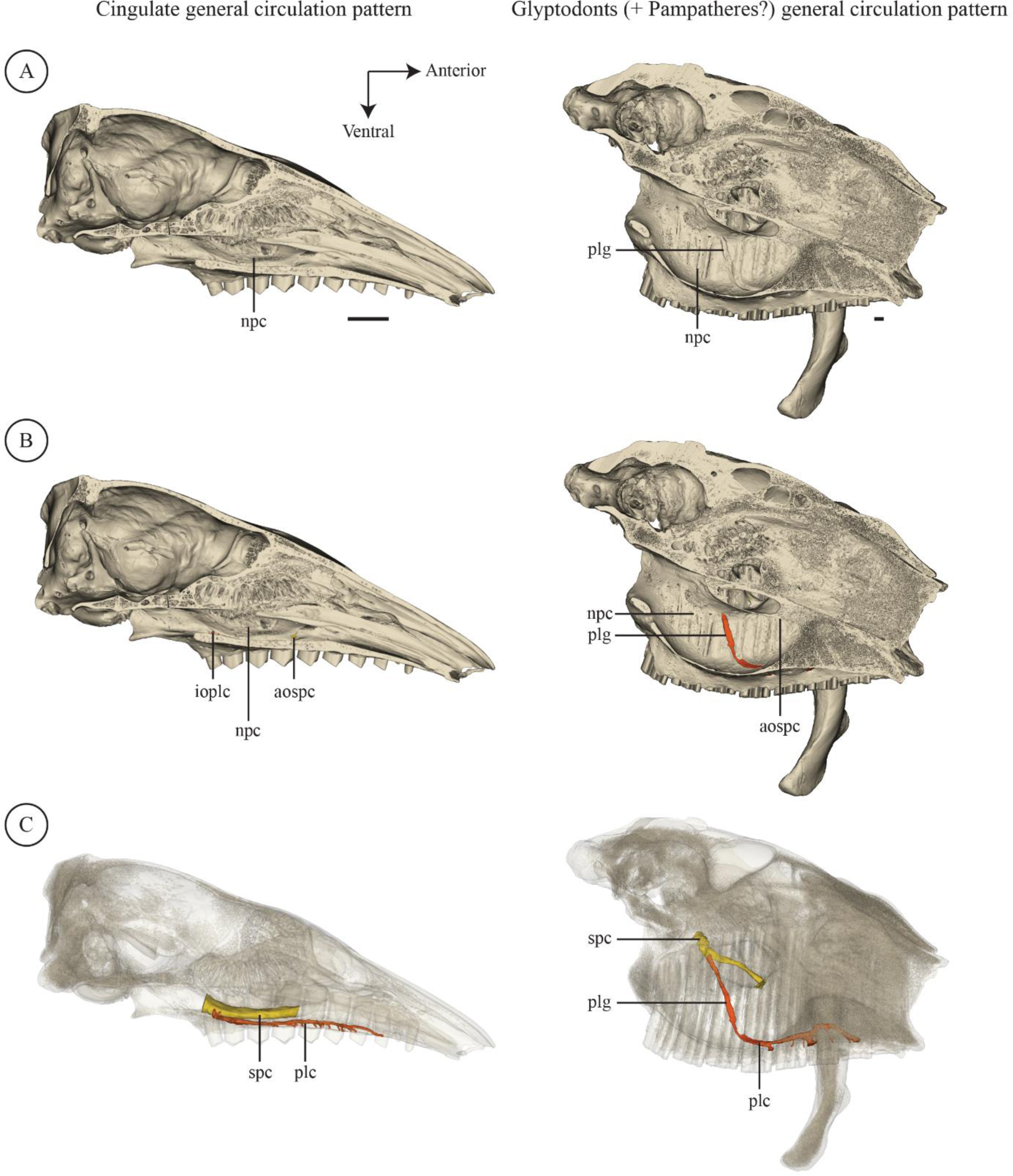
Comparative illustration of the trajectory of the palatine canal in glyptodonts and other cingulates (illustrated here by *Euphractus sexcinctus* (AMNH 133304)). A, lateral internal view on a cranium sectioned on the midline. B, same view as A but with the palatine canal reconstructed. C, same view as A with cranium transparency and with the sphenopalatine and palatine canals reconstructed. Abbreviations: aospc, anterior opening of the sphenopalatine canal; ioplc, internal opening of the palatine canal; npc, nasopharyngeal canal; plg, palatine grove; spc, sphenopalatine canal. Scale = 1cm.

**FIGURE S7.**
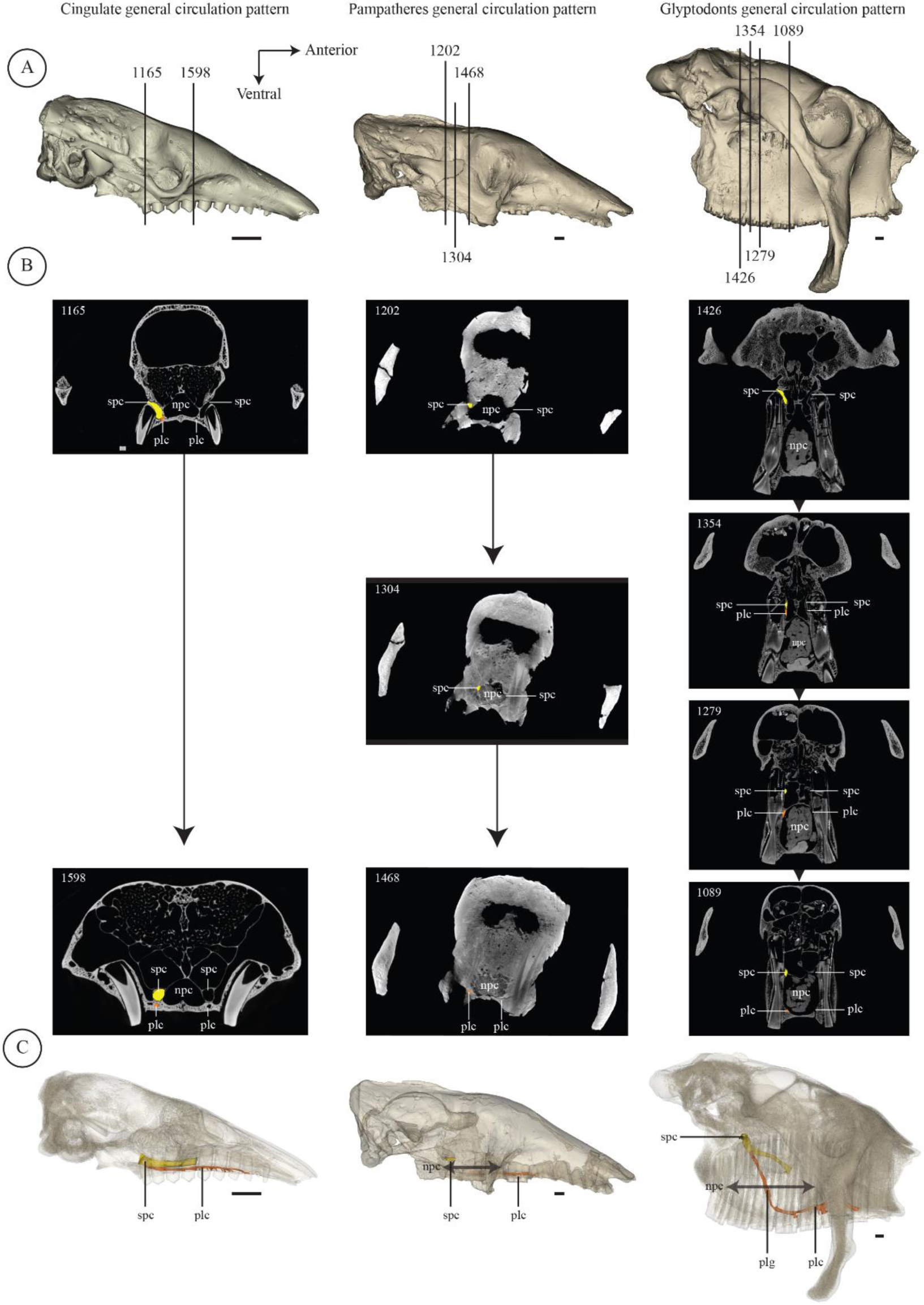
Illustration of the position of the sphenopalatine and palatine canals in relation to the nasopharyngeal canal in *Euphractus sexcinctus* (AMNH 133304), *Vassallia maxima* (FMNH P14424) and *Glyptodon* sp. (MNHN.F.PAM 760). A. Selected slides (numbered according to their position in the image stack) indicated on the cranium in lateral view. B. Slides in coronal view. C. Canals are indicated on cranium in transparency in lateral view (same as A). The same colors were used for canals: orange, palatine canal; yellow, sphenopalatine canal. Abbreviations: npc, nasopharyngeal canal; plc, palatine canal; plg, palatine groove; spc, sphenopalatine canal. Scale = 1 cm.

**FIGURE S8.**
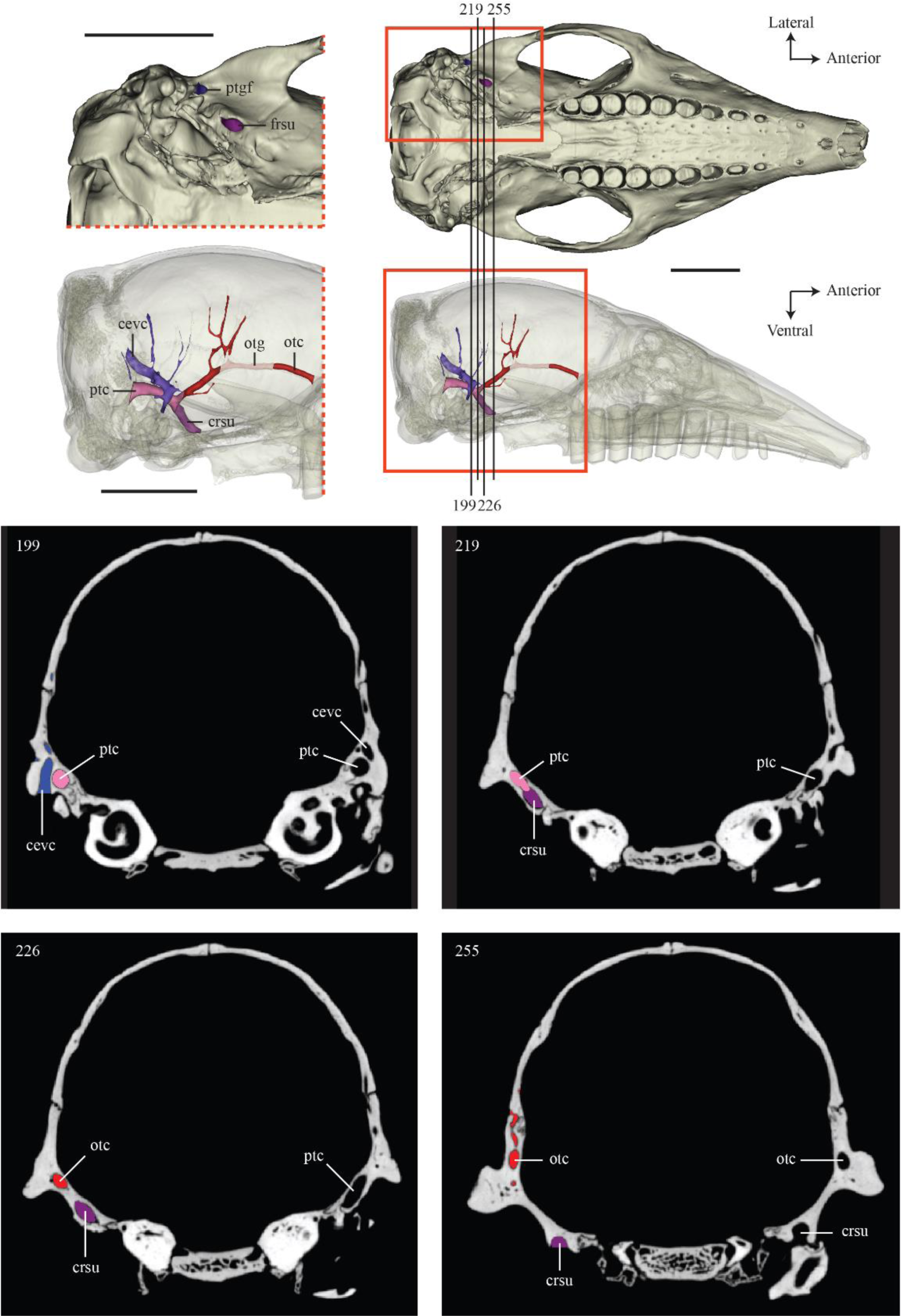
Identification of the canal for the ramus superior of the stapedial artery in *Tolypeutes matacus* (FMNH 28345). External opening is illustrated on a cranium in ventral view with a focus on the auditory region. Canal trajectory in relation to the other braincase canals is representing on a transparent cranium in lateral view with a focus on the neurocranium. Several parts of the canal (external opening, contact with the orbitotemporal and posttemporal canal, etc.) are also illustrated by numbered slides (in relation to the total number of slides) and indicated on the crania. Abbreviations: cevc, capsuloparietal emissary vein canal; crsu, canal for the ramus superior of the stapedial artery; otc, orbitotemporal canal; otg, orbitotemporal groove; ptc, posttemporal canal. Scale = 1 cm.

**FIGURE S9.**
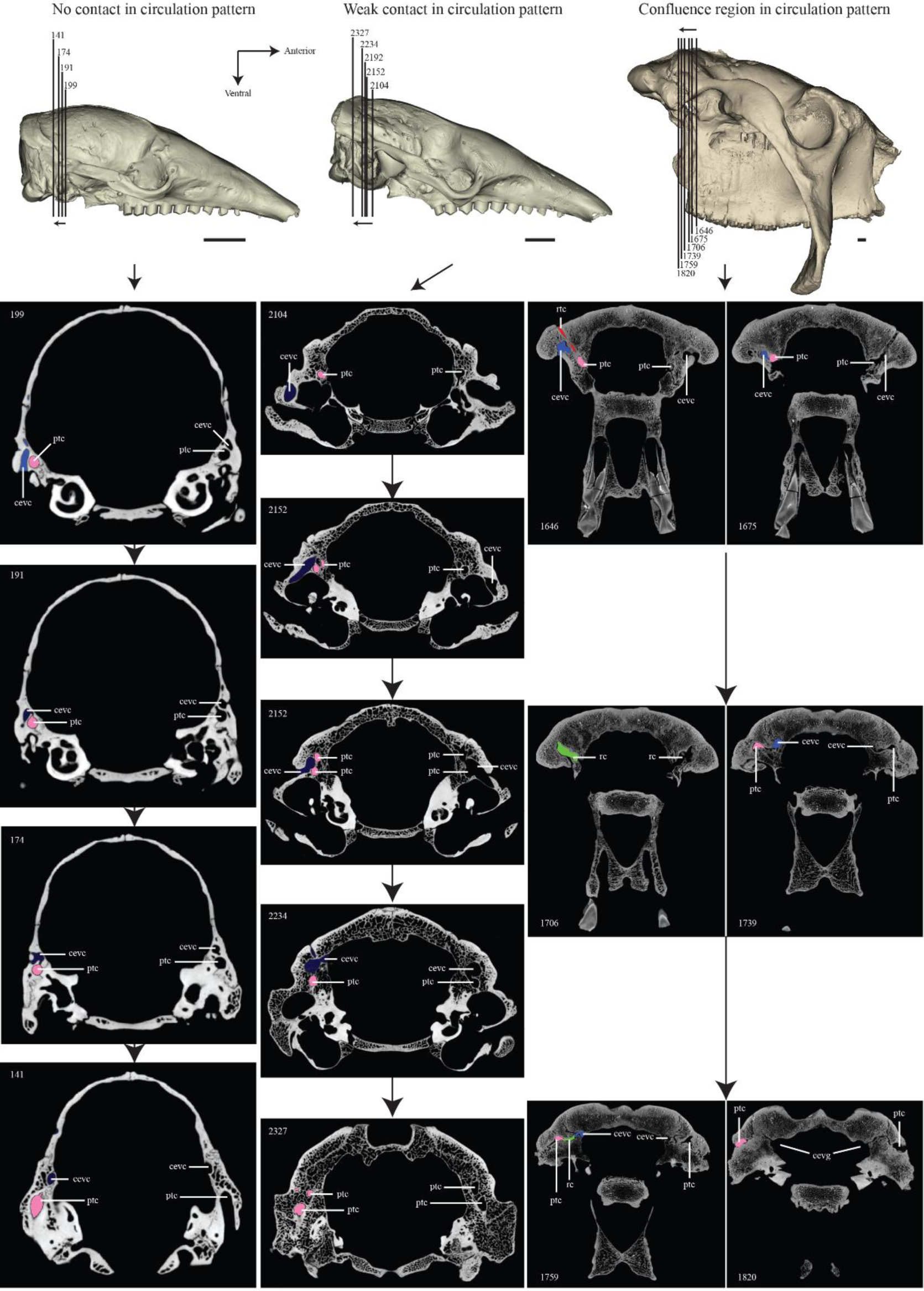
3 Different cases of contact/non-contact between the three braincase canals analyzed in our study (including the region of confluence) in *Tolypeutes matacus* (FMNH 28345), *Euphractus sexcinctus* (AMNH 133304) and *Glyptodon* sp. (MNHN.F.PAM 760). Selected slides (numbered according to the total slide number) are indicated on cranium in lateral view and canals are colored on one side of each slide. Abbreviations: cevc, capsuloparietal emissary vein canal; cevg, capsuloparietal emissary vein groove; ptc, posttemporal canal; rc, region of confluence; rtc, ramus temporalis canal. Scale = 1 cm.

